# Synthetic Serum Markers Enable Noninvasive Monitoring of Gene Expression in Primate Brains

**DOI:** 10.1101/2025.06.01.657212

**Authors:** Sangsin Lee, McKenna D. Romac, Sho Watanabe, Mykyta Chernov, Honghao Li, Emma Raisley, Jerzy O. Szablowski, Vincent D. Costa

## Abstract

We demonstrate a noninvasive approach to measure transgene expression in the brain of nonhuman primates using blood tests with engineered reporters called Released Markers of Activity (**RMAs**). RMAs can exit the brain and enter the bloodstream via reverse-transcytosis across the blood-brain barrier. We demonstrate that these reporters can be used to repeatedly monitor expression of multiple transgenes in cortical and subcortical brain regions simultaneously over a period of three months. RMAs are also sensitive enough to detect circuit-specific Cre-dependent AAV expression. Through this study, the RMA platform provides a cost-efficient, noninvasive tool for neuroscience study of large animals, enabling sensitive, multiplexed, and repeatable measurements of gene expression in the brain with a blood test.

## INTRODUCTION

The lack of a simple, inexpensive, and noninvasive method for repeated monitoring and quantification of *in vivo* transgene expression in the nonhuman primate (NHP) brain over long periods of time is a barrier to the use of molecular neuroscience tools in NHPs. While repeated *in vivo* neuroimaging of transgene expression using positron-emission tomography (PET)^1,2^ and magnetic resonance imaging (MRI)^4^ is possible in some cases, doing so requires repeated anesthesia events, specialized equipment, and the development of specialized contrast agents. Recent advances in noninvasive neuroengineering resulted in the development of genetically encoded contrast agents for various tissue-penetrant forms of energy, such as ultrasound^5^, radiofrequency waves^6,7^, and light^8^. While these tools have the potential to improve monitoring of transgenes, imaging transgene expression in the larger primate brain—especially within deep, subcortical structures^9^— still requires use of expensive equipment and anesthesia. An immediate need therefore exists for a simple scalable technology for long-term monitoring of transgene expression in NHPs with the ability to easily detect transgene expression activities regardless of where in the brain a gene is expressed.

Released Markers of Activity, or RMAs, were developed to address these limitations and enable monitoring of transgene expression in genetically-labeled cell populations with a simple blood test^10^. RMAs are genetically-encoded synthetic serum markers that are produced in the brain, but can be transported into the bloodstream, where they can be detected using simple blood tests^10^. We previously successfully demonstrated this versatile platform in rodent models, achieving sensitivity in detecting as few as 12 neurons and tracking changes in neuronal activity through genetic tagging of the immediate early gene *Fos* with RMAs^10,11^. We have also shown that RMAs can be used to monitor transcripts through the use of RNA-based sensors^12–14^.

RMAs consist of three components (**Fig. 1A**). First, they contain a signaling secretion sequence to enable secretion from the cells into the interstitial space. To make RMAs detection simple and sensitive we fused this sequence to an easily detectable reporter, a luciferase. Finally, that assembly was fused to the Fc region of immunoglobulin G (IgG) antibody which enables crossing through the intact blood-brain barrier (BBB) into the blood using neonatal Fc receptor (FcRn)-mediated reverse transcytosis^15^ (**Fig. 1B**). Additionally, the Fc domain added to the RMA prolonged the serum half-life^16,17^ increasing the detection sensitivity, leading to successful detection of transduction in as few as 12 neurons in the mouse brain with 5 microliters of serum. The RMA originally designed for rodents includes the naturally secreted enzyme *Gaussia* luciferase (Gluc) and the mouse Fc region^10^.

**Figure 1.**
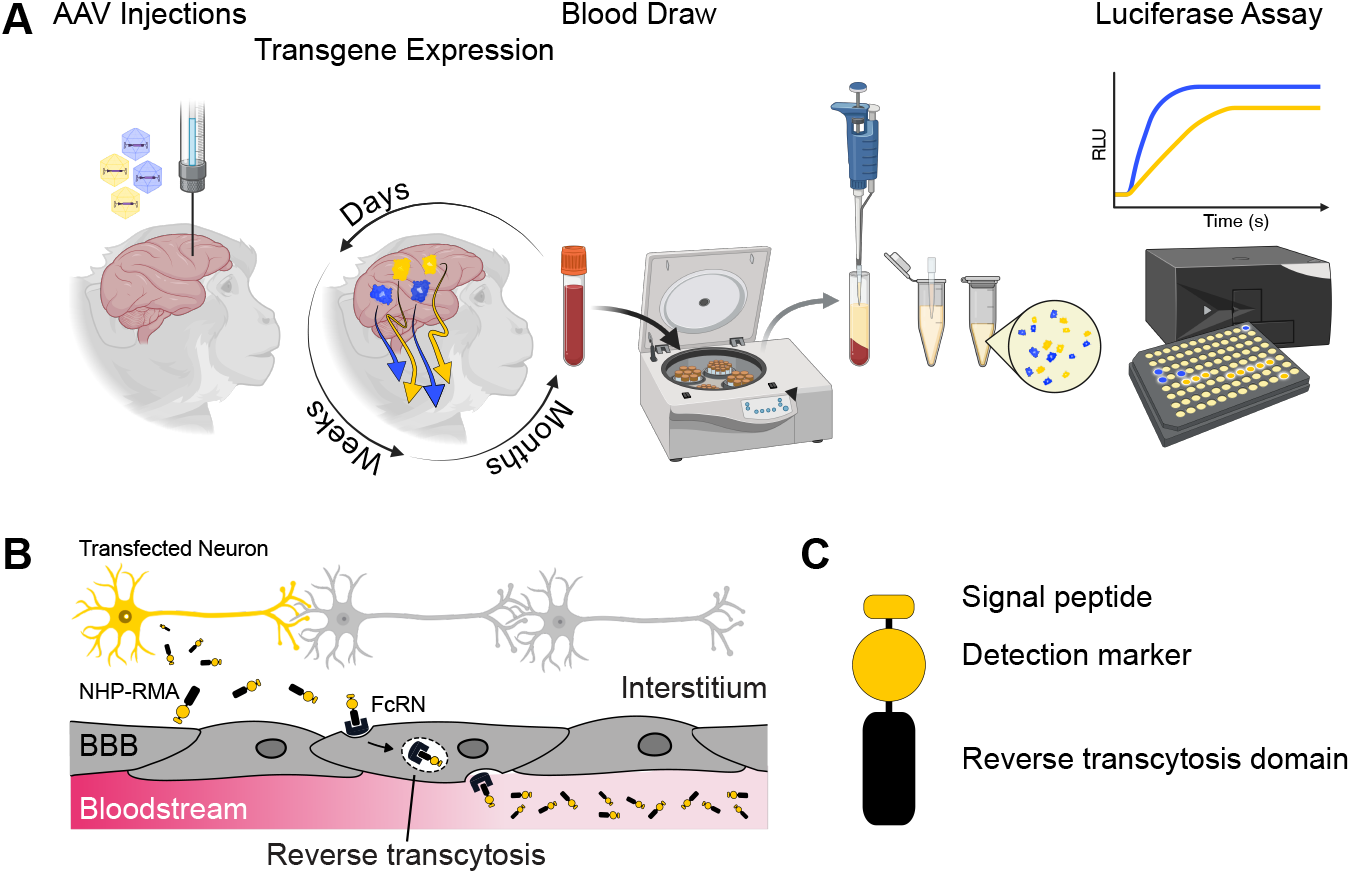
Noninvasive monitoring of transgene expression in macaque brains using NHP-RMAs. **A) Experimental workflow for using NHP-RMAs.** NHP-RMAs are first injected into a target brain area, then repeated blood draws are collected to monitor transgene expression, followed by serum processing preparation of luciferase assays that detect and quantify the presence of the NHP-RMAs in the blood. These released reporters from the collected blood can be measured using any compatible biochemical detection methods, avoiding the need for direct brain tissue imaging and enabling sensitive, long-term monitoring of gene expression in individual subjects. **B) Mechanism of NHP-RMA transport from brain to blood**. A genetically labeled neuron (yellow) expresses and secretes NHP-RMAs into the surrounding tissue. After entering the interstitial space, the NHP-RMAs bind to the targeted FcRn receptors, which mediates reverse transcytosis across the BBB. The NHPRMAs dissociate from the FcRn receptor upon reaching the bloodstream and are released into circulation. **C) NHP-RMA protein design**. The reporter consists of three components:1) an N-terminal secretion signal peptide, 2) a detectable protein marker, (e.g. *Gaussia* or *Cypridina* luciferase) which can be substituted with a fluorescent protein or an antibody epitope depending on the readout method, and 3) the Fc domain from the heavy chain region of a macaque antibody to facilitate BBB crossing via reverse transcytosis.

To adapt RMAs for use in macaques, we swapped the mouse IgG1 Fc region into its allogeneic variant in macaques. We retained the Gluc domain, which already contains the mammalian secretion signal and was proven as a sensitive luciferase marker for detection via bioluminescence imaging (**Fig. 1C**). We found that the NHP variant of RMAs (NHP-RMAs) was capable of crossing through the BBB after intraparenchymal brain injection, and could report on the AAV-based gene delivery. Additionally, we show evidence of monitoring Cre-based recombination in the intact NHP brain through multiple blood tests. No other noninvasive gene expression monitoring modality currently is comparable to RMAs for high sensitivity, high throughput, repeated monitoring of multiple transgene expression activities in the mammalian brain, regardless of its size.

## RESULTS

### Development of an NHP variant of RMAs (NHP-RMA)

We surveyed the macaque Fc domains of different IgG antibody subclasses (IgG1 to IgG4) and aligned their sequences with that of the mouse IgG1 Fc used in the original RMA study (**Fig. 2A**). The alignment revealed no sequence gap and a high conservation of physicochemical properties across Fc residues in different macaque IgG subclasses and the mouse Fc. Furthermore, the histidine residues located in the Fc hinge region (denoted by red triangles in Fig. 2A) were fixed across all Fc sequences. These histidine residues confer the pH-dependent binding function^15^ of the Fc receptor, suggesting both intra- and inter-species conservation of the reverse transcytosis mechanism that allows traversing of the BBB. For our final design we selected the macaque IgG1 Fc to match the host species and fused it with the Gluc domain (**Fig. 1C**) and referred to the designed protein as NHP-RMA.

**Figure 2.**
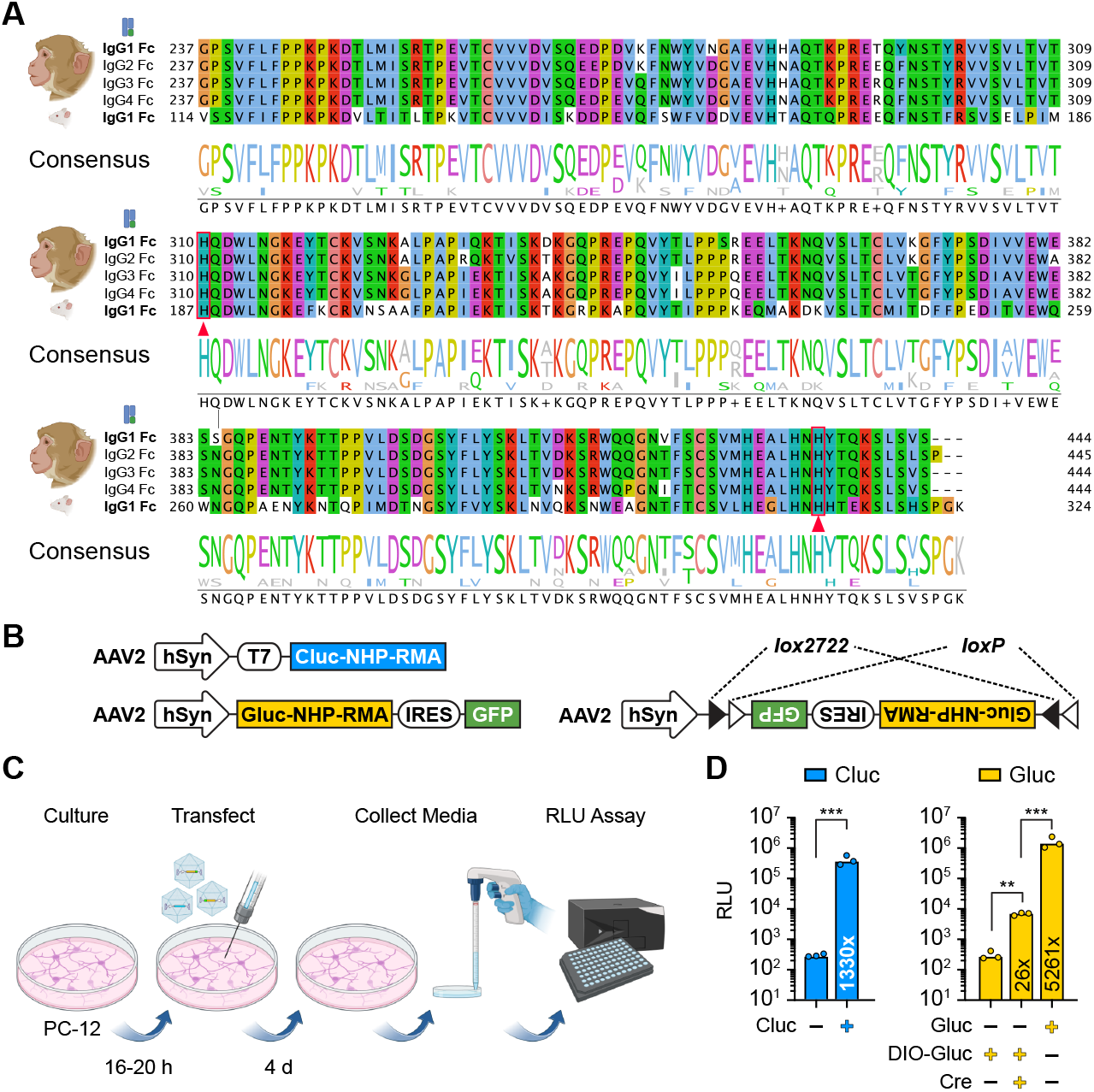
Bioengineering of NHP-RMAs. A) Sequence alignment of the Fc domains of different IgG subclasses. For macaque IgG1 to IgG4, Fc sequences were obtained from the PDB IDs 6D4E, 6D4I, 6D4M, and 6D4N, respectively. The mouse IgG1 Fc sequence was included to evaluate conservation between macaque and mouse species. Red triangles mark the H310 and H435 histidine residues critical for pHdependent binding to FcRn. Conservation scores were calculated based on the similarity of physicochemical properties, with score of 10 indicating complete conservation (see Methods). The consensus sequence shows the frequency of amino acid at each residue position. **B) Vector diagrams of the AAVs created to express NHPRMAs in the nonhuman primate brain**. Three AAVs were designed to express NHP-RMAs based on either *Cypridina* or *Gaussia* luciferase in neurons using the *hSyn* promoter. One of the AAVs was Cre-dependent. **C) Experimental workflow for *in vitro* functional validation of the AAVs in rat PC-12 cells**. Neurons were cultured and then transfected with specific AAVs (AAV dose with a multiplicity of infection of 100,000), after which the tissue media was sampled. Luciferase assays of the tissue media were then used to detect the NHP-RMA protein of interest. **D) Bioluminescence signals measured from PC-12 cell culture media containing secreted NHP-RMAs**. Increased bioluminesence was detected in the tissue media following AAV transfection of neurons to directly produce Cluc- and Gluc-based NHP-RMA proteins. There was no change in the bioluminesence of the tissue media following AAV transfection of neurons with the DIO-Gluc based NHP-RMA until the same cells were transfected with an AAV that expressed Cre-recombinase. Numbers within bars represent fold increases in the bioluminesence signal (RLU) compared to the no-expression control: Cluc (*n*=3 independent cultures) and Gluc/DIO-Gluc (*n* = 6) analyzed. + and – symbols indicate when a particular AAV was added to the tissue culture. Data are shown as mean ± SD.

The primary benefit of using NHP-RMAs to monitor transgene expression in the brain is the ability to repeatedly monitor gene expression over weeks to months using repeated blood draws from the same monkey (**Fig. 1A)**. We constructed three adeno-associated viral (AAV) vectors encoding NHP-RMAs under the neuron-specific *hSyn* promoter (**Fig. 2B**) for the purpose of tracking transgene expression *in vivo*. Two of the AAV-produced NHP-RMAs were Gluc-based. By replacing the Gluc domain with an orthogonal luciferase, we also made a third NHP-RMA using *Cypridina* luciferase (Cluc) as a reporter to enable multiplexed monitoring of transgene expression (**Fig. 2B**). The functionality of all three AAVs was first validated by transducing PC-12 cells (a common rat cell line used for neurosecretion studies)^18^ followed by detecting release of the NHP-RMAs in the culture media (**Fig. 2C**). We detected more than a 5000-fold increase in the media of cells transfected with AAV2-hSyn-Gluc-IRES-GFP (*t*(2) =4.084, *p* = 0.015) and more than a 1000-fold increase in the media of cells transfected with AAV2-hSyn-Cluc (*t*(2) =4.798, *p* = 0.0087), compared to samples of control media from uninfected cells. We also detected more than a 25-fold increase in the media of cells that were co-transfected with AAV2-hSyn-DIO-Gluc-IRES-GFP and AAV2-hSyn-CRE-GFP (*t*(2) = 11.897, *p* < 0.001; **Fig. 2D**).

### Reverse transcytosis and detection of NHP-RMA protein *in vivo*

The tissue culture results validated the functionality of the AAVs and experimental workflow for measuring the NHP-RMA signal. We verified whether the NHP-RMA protein produced by neuronal expression of the transgene would be bound by the targeted Fc receptor and undergo reverse transcytosis into the bloodstream and remain stable within the circulation. We produced a purified version of the NHP-RMA protein (**Fig. 3A and Fig. S1**). Given that the Fc domain is known to facilitate endosomal recycling of proteins and extend half-lives in the circulatory system^15^, we first examined the pharmacokinetics of NHP-RMAs (**Fig. 3B**) by intravenously (IV) administering purified proteins and measuring the rate of signal decay overtime. Testing a large and small dose revealed the *β*-phase half-life (*t1/2*) ranged between 1.2-1.5 days (**Fig. 3C)**, which is longer than the 20-minute half-life of Gluc alone^19^ but shorter than the 4-day half-life of a Gluc-RMA previously used in mice^10^. To validate that our detected signals represented the intact activity of NHP-RMAs in serum we directly mixed different concentrations of the NHP-RMA purified protein with serum samples from mice. We confirmed a strong positive, linear correlation between bioluminescence signals and protein concentrations, consistent with the known characteristics of Gluc^20^ (**Fig. S2A**). Additionally, multiple freeze-thaw cycles of the spiked serum samples did not degrade NHP-RMA luciferase activity (**Fig. S2B**).

**Figure 3.**
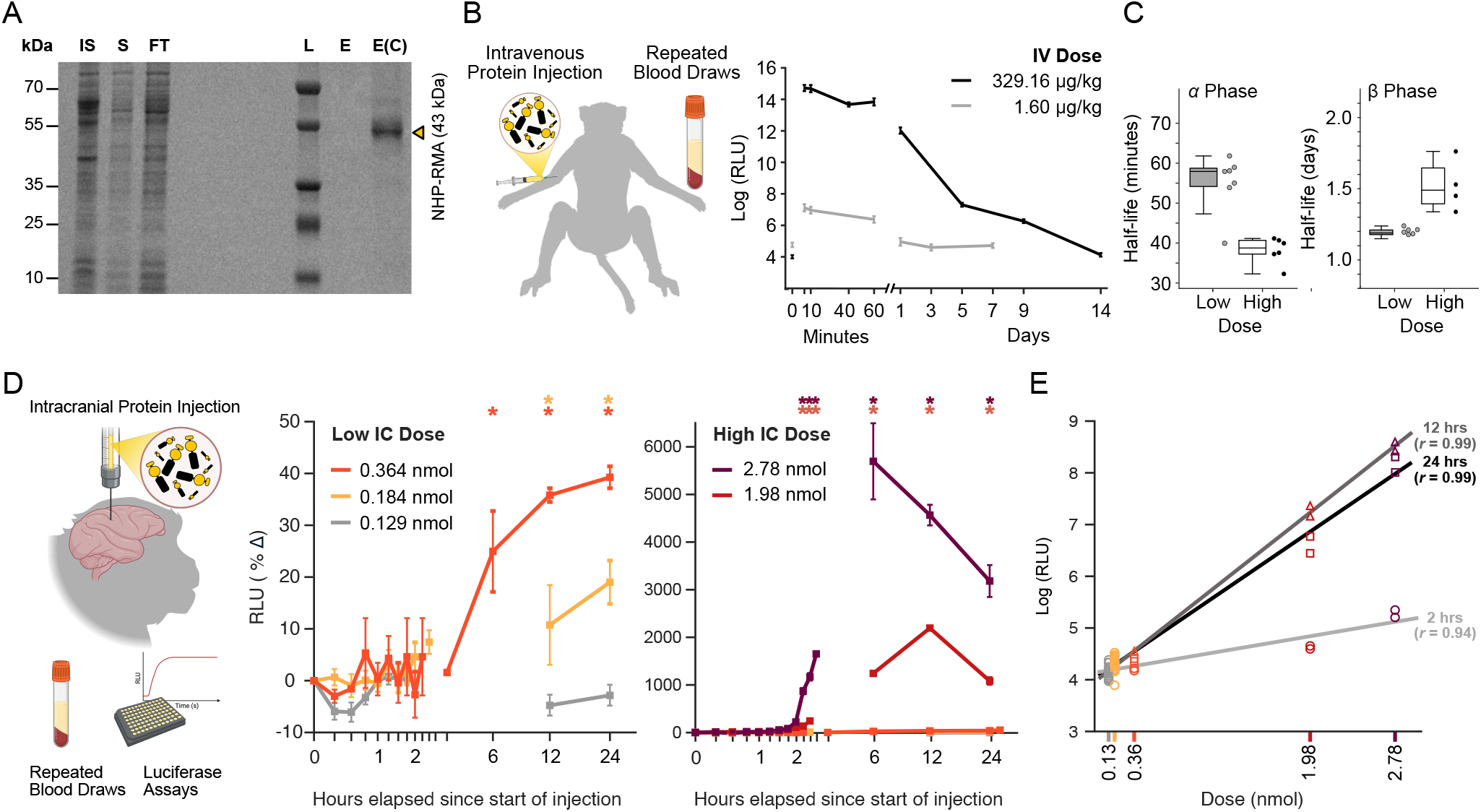
Direct injections of NHP-RMA protein into the brain undergoes reverse transcytosis into the bloodstream. **A) Purification of NHP-RMA protein.** SDS-PAGE analysis of NHP-RMA proteins isolated from bacterial expression. (IS – insoluble fraction, S – soluble fraction, FT – flow-through, W – wash, E – elution). **B) Bioluminescence signals following intravenous injections of the purified NHP-RMA protein into the bloodstream**. IV injections of the purified protein were followed up with blood draws at 5, 10, 40, and 60 minutes and at intermittent timepoints from 1 to 14 days after the injection. Mean ± SEM of the Gluc based luciferase assay for two monkeys that received a low or high dose of the purified protein. There was an immediate increase in signal after 5 minutes that declined within an hour of the injection and continued to decline over the next 1-2 weeks. **C) Pharmacokinetics for clearance of the NHP-RMA protein from the blood supply following an IV injection**. Two compartment elimination model for *α*- and *β*-phase half-lives was used to analyze the pharmacokinetic parameters. Technical replicates (*n* = 3) were measured for each blood sample and the half-lives were estimated for each series of technical replicates. Data are shown as mean ± SEM. **D)** Intracranial injections of the purified protein were followed up with blood draws every 15 minutes after the start of the injection up to 3.25 hours after the injection. Additional blood draws were collected between 6 and 24 hours after the start of the injection. Mean ± SEM of the Gluc based luciferase assay for the monkeys that received injections into the putamen of different doses of the NHP-RMA protein. Colored asterisks above each timepoint indicate a significant increase (*p* < .05) in the RMA signal above baseline for specific doses. **E) Dose dependent increases in RMA signal following intracranial viral injections**. Linear regression indicated a positive correlation between the amount of protein injected into the brain and the RMA signal was observed as early as two hours after the protein was injected (*r* > 0.87, p < 0.05). Each point is a technical replicate for injected dose and timepoint: circle (2 hours), squares (12 hours), triangles (24 h).

Having confirmed that the half-life of NHP-RMA protein was longer than 1 day, we next determined the efficacy of the NHP-RMA protein in crossing the BBB after intracranial injections into the brain. Proteins with over 95% purity **(Fig. S1)** were injected to minimize brain toxicity, as contaminants from bacterial purification could trigger an inflammatory response that increases BBB permeability^21^ and allow NHP-RMAs to exit the brain through unintended pathways other than reverse transcytosis. All intracranial injections targeted the putamen, either bilateral (for 0.129 nmol, 1.98 nmol and 2.76 nmol doses) or unilateral for all other doses (0.184, 0.364). Because the NHP-RMAs—if effective— would allow us to confirm a successful injection and we were uncertain about whether gadolinium contrast agent (GAD) might interact with the NHP-RMA protein we did not collect post-surgery MRI scans. We collected serum samples immediately prior to starting each injection. We then sampled 1.5 mL of blood every 15 minutes up until 3.25 hours after the start of the protein injections, with additional blood draws acquired at 6, 12, and 24 hours after the start of the protein injection (Fig. 3D). At the lowest dose (0.129 nmol) we did not observe a change from baseline at 12 or 24 hours (Time: *F*(2,14) = 3.09, *p* = 0.077). At the next highest dose (0.184 nmol) we detected a 1.19 fold change at 24 hours that differed from baseline (*t*(14).= 3.07, *p* = 0.0082). However, this effect was highly dependent on the number of technical replicates assayed. When we doubled the amount of protein injected (0.364 nmol) we readily detected a 1.35- and 1.39-fold increase in the RMA signal at 12 and 24 hours relative to baseline (Time: *F*(2,2) = 156.7, *p* < 0.0001) and using fewer technical replicates (*n* = 2). Injecting substantially larger doses (1.98 and 2.76 nmol) led us to detect larger (11.19-to 56.95-fold) changes in the RMA signal relative to baseline between 6 and 24 hours after the injections (1.98 nmol: Time, F(9,9) = 710.6, *p* < 0.0001; 2.76 nmol: Time, F(9,9) = 196.54, *p* < 0.0001). The RMA had returned to baseline when blood draws were taken a minimum of 3 weeks after the injection procedure (for all comparisons *p* > 0.35). When we assessed dose-dependent changes in the RMA signal at each timepoint, a stable linear dose response emerged as early as two hours (critical *r* value > 0.811, *p* < 0.05; **Fig. 3E**) after starting the injections and there was a strong, positive correlation between the Gluc-based signal and the amount of protein injected at 12 and 24 hours after starting the injection—just as we saw when we spiked mouse serum with specific quantities of the purified protein (**Fig. S2A**). Overall, these results establish that the NHP-RMA protein crossed the BBB as expected and provide a reference frame for determining the amount of secreted protein in the brain following transgene expression of Gluc-based RMAs in neurons. It also highlights the potential use of co-injecting the NHP-RMA protein with other AAVs to verify whether an injection was successful in scenarios where co-injecting with contrast agents and post-surgical MRIs are not feasible.

### AAV-mediated gene expression in neurons to secrete NHP-RMA proteins in nonhuman primates

Use of AAVs to facilitate neuronal secretion of RMA proteins was previously demonstrated in mice^10^. Considering the genetic overlap in the mouse and macaque IgG1 Fc receptors and our *in vitro* validation of the designed AAVs (**Fig. 2**), we conducted parallel evaluations of the AAVs designed to express NHP-RMA proteins in wild-type mice and rhesus macaques.

We co-injected the AAV2-hSyn-Gluc-IRES-GFP (125 nl) and AAV2-hSyn-Cluc (125 nL) into the basal ganglia of the mice and evaluated RMA signals just prior to the injection and after 1 and 2 weeks of expression (**Fig. 4A**). The stock titers are provided in the methods section. Histological quantification using in situ hybridization to detect Cluc (*F*(1,2) = 15.47, *p* = 0.05) expression in neurons and immunofluorescence to detect Gluc (*F*(1,2) = 27.69, *p* = 0.0343) and associated GFP (*F*(1,2) = 50.68, *p* = 0.0127) expression in neurons confirmed expression of the NHP-RMA transgenes was restricted to the injected hemisphere (Hemisphere: *F*(1,2) = 203.49, *p* = 0.0049; Hemisphere x Marker Protein: *F*(2,4) = 3.38, *p* = 0.1404). Of the sampled nuclei, 15.86% expressed Gluc and 16.5% expressed Cluc (**Fig. 4B**). Consistent with the *in vitro* experiments in cultured rat neurons, we observed an increase in the bioluminescence signal associated with each RMA (**Fig. 4C**). In mice, the Cluc-based RMA signal increased 1009-fold after 2 weeks (Time: *F*(2,4) = 12.23, *p* = 0.0198) and the Gluc-based RMA signal increased 1,928-fold after 2 weeks (Time: F(2,4) = 361.27, p < 0.0001). At 2 weeks, the fold increase in the Gluc-based signal was larger than increase in the Cluc-based signal (Luciferase x Time: *F*(2,4) = 11.28, *p* = 0.0227).

**Figure 4.**
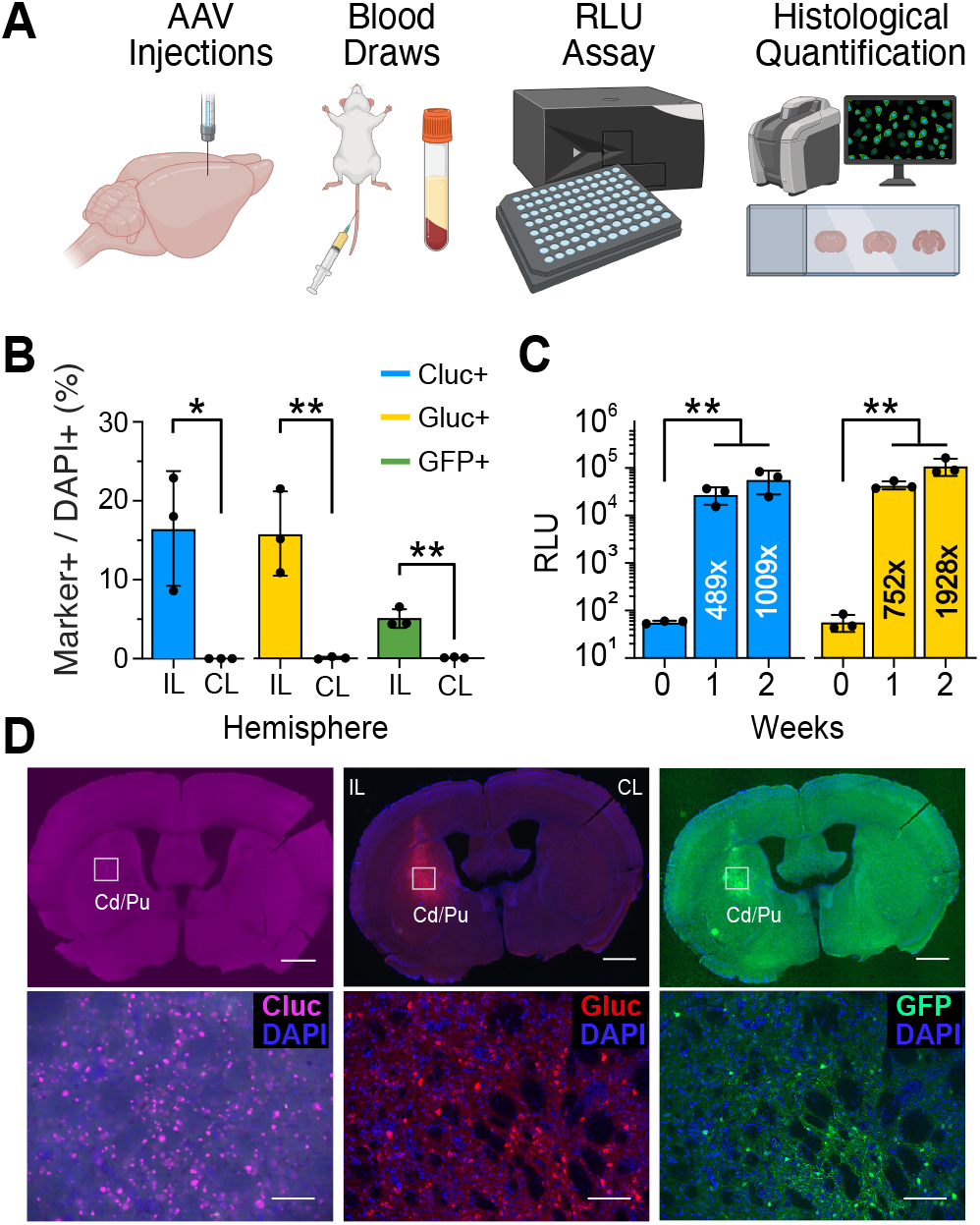
Cross-species validation of AAV mediated gene expression of NHP-RMAs. **A) Experimental workflow for in vivo functional validation of the designed AAVs in mice.** AAVs encoding Gluc- and Cluc-based RMAs were co-injected in the rodent basal ganglia and their expression was measured through luciferase assays of collected blood samples. Genetic overlap in the Fc receptor suggests that the designed AAVs should mediate NHP-RMA neuronal protein secretion in mice. **B) Histological quantification of NHP-RMA markers in mouse basal ganglia**. Cluc+ cells were quantified using in situ hybridization as the AAV lacked a fluorescent reporter and there were no effective primary antibodies validated for macaque tissue. Gluc+ and GFP+ cells were detected using immunofluorescence. Cell counts were normalized against DAPI+ nuclei detected within the imaged region of interest in the hemisphere ipsilateral (IL) to the injection or contralateral (CL) within the same tissue section. Datapoints represent cell counts for individual mice. **C) Serum bioluminescence signals for Gluc and Cluc**. Numbers shown inside each bar indicate signal fold changes relative to the corresponding baseline (0 weeks). Data points represent luciferase assays for individual mice. **D) Examples of Cluc and Gluc expression**. Overview (4X) and detailed (20X) tissue sections depicting expression of Cluc RNA (left panel) and co-expression of Gluc (center panel) and GFP (right panel) proteins in two tissue sections through the basal ganglia of the same mouse injected with the AAVs. Scale bar, 1 mm (4X overview) and 100 um (20X high-magnification). Cd = caudate; Pu = putamen; *\* p* < 0.05; *\*\* p* < 0.01. All data are shown as mean ± S.D.

We injected the same two AAVs into the brains of five rhesus macaques to transfect neurons and enable secretion of Cluc- and Gluc-based NHP-RMA proteins (**Fig. 5A**). We injected AAV2-hSyn-Cluc into the basal ganglia (putamen or nucleus accumbens) of all five monkeys and in two of the monkeys we also injected it at multiple sites in the prefrontal cortex (**Table S3**). We injected AAV2-hSyn-Gluc-IRES-GFP into the basal ganglia of three of the monkeys and in one of those monkeys we also injected it at multiple sites in the anterior cingulate and frontopolar cortex. All the injections were at lower titers than those used in mice (**Table S3**). We collected a blood sample just before we injected the AAVs. To monitor AAV-mediated transgene expression, blood was drawn 24 hours after starting the injection and then once a week for up to 12 weeks **(Table S3)**. Luciferase assays were then used to quantify changes in the Gluc- and Cluc-based signals (**Fig. 5A**) by generating 3-8 technical replicates of each timeseries for each monkey.

**Figure 5.**
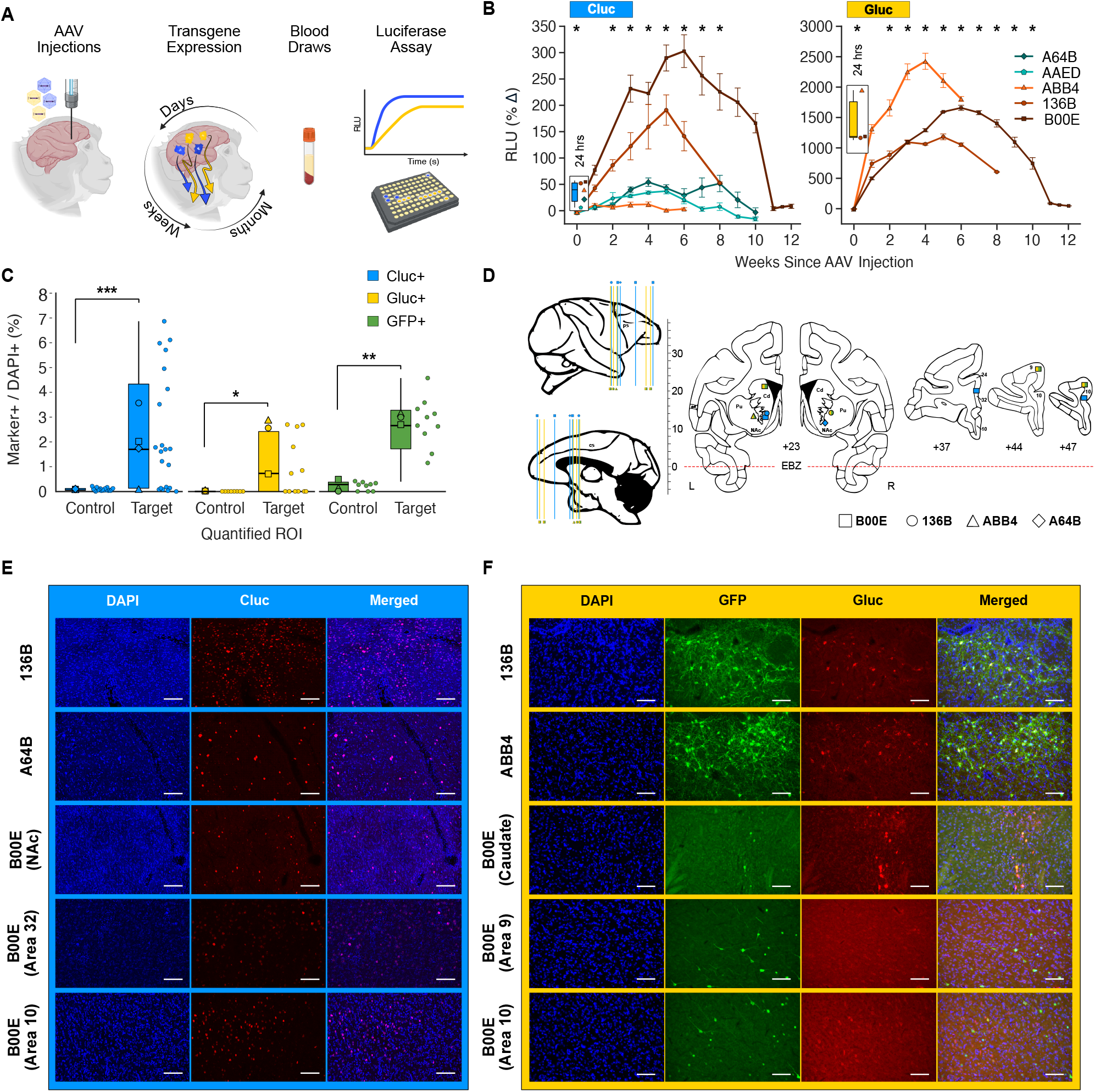
AAV-mediated neuronal secretion of Gluc- and Cluc-based NHP-RMA proteins in monkeys. **A) Experimental workflow for *in vivo* functional validation of the designed AAVs in monkeys.** AAVs encoding Gluc- and Cluc-based RMAs were either co-injected into the same brain region or injected into separate brain regions in rhesus macaques (see Table S3 for details). Blood draws were taken at different timepoints to track transgene expression through luciferase assays to readout RMA signal levels. **B) Time course of RMA signals following AAV injections in individual monkeys**. Mean ± SEM across technical replicates of the time courses of the Cluc- and Gluc-based RMA signal change relative to baseline. Asterisks above each timepoint indicate an increase (*p* < .05 Bonferroni corrected) in the RMA signal above baseline based on a generalized linear mixed model that assessed time-varying changes, averaged across all monkeys. Inset panels depict boxplots of the distribution of RMA signal change relative to baseline 24 hours after the AAV injections. Datapoints represent estimates for each monkey. **C) Histological quantification of NHP-RMA markers in the macaque brain**. Cluc+ cells were detected using in situ hybridization and Gluc+ and GFP+ cells were detected using immunofluorescence. Boxplots show the distribution of cell counts within regions of interest surrounding the injection site and or adjacent control regions pooled across monkeys. Individual symbols represent mean estimates for each monkey. Cell counts were normalized against DAPI+ nuclei detected within the imaged ROIs. **D) Schematic showing location of representative tissue sections**. Locations of the representative sections shown in the bottom panels are depicted on illustrative sections from Saleem and Logothetis (2012)^3^.… (contd…) Shapes indicate from which monkey the tissue section came and the color of the shape indicates whether the section was used to test for expression of Cluc (blue) or Gluc/ GFP (yellow/green). **E) Representative in situ hybridization images of Cluc expression in the basal ganglia and prefrontal cortex**. High-magnification (20X) images of Cluc (red) expression in two monkeys within the ventral striatum (vSTR; first two rows) or nucleus accumbens (NAc; third row), anterior cingulate cortex (Area 32; fourth row) and frontopolar cortex (Area 10; fifth row) in a third monkey. **F) Representative immunofluorescence images of Gluc and GFP expression in the basal ganglia and prefrontal cortex**. High-magnification (20X) images of Gluc (red) and GFP (green) expression in the putamen (Pu) of two monkeys (first and second row), and in the caudate nucleus (Cd; third row), anterior prefrontal cortex (Area 9; fourth row), and frontopolar cortex (Area 10; fifth row) in a third monkey. Ps = principal sulcus; Cs = cingulate sulcus; EBZ = ear bar zero.

We detected increases in both the Cluc- and Glucbased signals compared to baseline 24 hours after injecting the AAVs (**Fig. 5B**). There was a 1.35-fold (±0.0315 SEM) increase in the Cluc signal compared to baseline at 24 hours (*t*(25) = 11.15, *p* < 0.0001) and this effect varied across the five monkeys (Monkey: F(4,24) = 8.74, p = 0.0002; range: 1.07-fold to 1.55-fold). There was a larger, 15.33-fold (±0.44 SEM), increase in the Gluc signal compared to baseline at 24 hours (*t*(9) = 32, *p* < 0.0001) and this effect also varied by monkey (*F*(2,9) = 30.78, *p* < 0.0001). This was surprising but not implausible considering that AAV dsDNA synthesis can occur within hours after AAVs transfect a cell^22^. One week later, however, both the Cluc-(*t*(24) = −2.36, *p* = 0.0266) and Gluc-based (*t*(9) = −13.09, *p* < 0.0001) signals had decreased relative to the 24-hour assay.

We observed a progressive increase in both the Cluc (Time: *F*(10, 179) = 11.91, *p* < 0.0001) and Gluc (Time: *F*(10, 70) = 15.87, *p* < 0.0001) signals up to 7 weeks after the AAV injections after which the signals started to decrease back towards baseline. The Cluc signal increased compared to the baseline measurement from two until eight weeks after the injection and peaked with a 2.1-fold increase after five weeks. The Gluc signal increased compared to the baseline from two until ten weeks after the injection and peaked with an 18.01-fold increase after 7 weeks. This pattern was observed in each monkey that we injected (**Fig. 5B**).

We randomly selected tissue sections from each monkeys’ brain surrounding the stereotaxic coordinates where we injected each of the AAVs (**Fig. 5C**) and specified target and control regions of interest (ROI). Using in situ hybridization to detect Cluc (*t*(27) = 5.09, *p* < 0.0001) we confirmed heightened expression in neurons within injected compared to control ROIs (**Fig. 5D and E**). Using immunofluorescence we confirmed heightened expression of Gluc (*t*(3) = 5.10, *p* = 0.0169) and associated GFP (*t*(5) = 5.88, p = 0.002) within injected (Gluc: M = 2.13%; GFP: M = 2.75%) compared to control ROIs (**Fig. 5D and F**).

### NHP-RMAs enable monitoring of circuit-specific Cre-dependent gene expression

Unlike in mice, developing transgenic nonhuman primates presents technical and logistical challenges. Therefore, circuit-specific chemogenetic and optogenetic perturbations in nonhuman primates require intersectional viral approaches for targeted gene expression in specific neuron types. We have demonstrated that Cluc- and Gluc-based NHP-RMAs could simultaneously monitor gene expression in different brain regions. We next sought to use a Cre-dependent version of the Gluc-based NHP-RMAs (**Fig. 6A**) to determine if we could temporally monitor Cre-Lox recombination-dependent expression of a transgene in a specific neural circuit.

**Figure 6.**
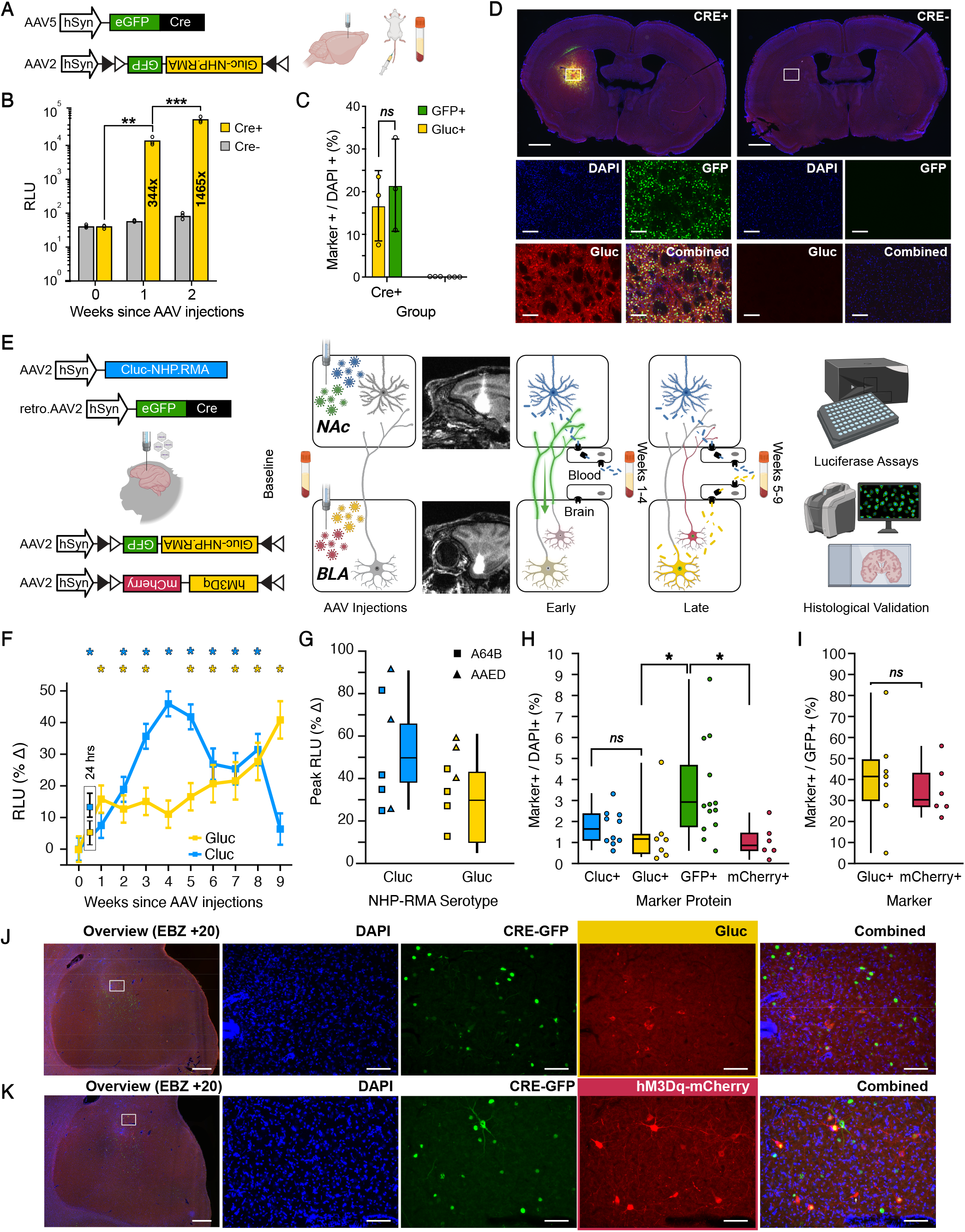
Monitoring gene expression in amygdalar circuitry using NHP-RMAs. **A) Experimental workflow for *in vivo* functional validation of the Cre-dependent Gluc-based RMAs in mice.** Schematic of viral vectors carrying Cre-dependent, double-floxed Gluc-NHP-RMA sequence for selective gene expression. The DIO-Gluc vector was either co-injected with the Cre vector or in isolation. … (contd…) Blood draws were taken at the time of injection and for two weeks following the AAV injections. **B) Bioluminescence signals measured from blood draws of mice injected with AAVs encoding Gluc NHP-RMA, with or without co-injection of Cre recombinase**. Bars represent mean Gluc-based signal at each timepoint in each group. Datapoints represent signal estimates for each mouse in each group. Numbers shown inside each bar indicate fold increases in signal relative to the corresponding baseline (0 weeks). **C) Credependent expression of NHP-RMA protein markers in mouse basal ganglia**. Gluc+ and GFP+ cells were detected using immunofluorescence. Cell counts were normalized against DAPI+ nuclei detected within imaged ROIs in the Cre+ and Cregroups. **D) Representative overview and high-magnification images of Cre-dependent Gluc expression in the presence of Cre recombinase**. NHP-RMAs are expressed in the tissue section on the left, which is from a mouse injected with Cre that exhibits GFP expression, while the tissue section on the right is from a mouse not injected with Cre and that shows no GFP expression and by extension no NHP-RMA expression. White rectangles mark the regions shown in the insets below the whole-brain images. Scale bars: 1000 µm (whole mouse brain) and 200 µm (high-magnification images). **E) Experimental workflow for using NHP-RMAs to monitor Cre-Lox recombination dependent transgene expression in amygdala circuitry**. Schematic of NHP-RMA viral vectors co-injected into either the NAc or BLA. The Cluc vector was co-injected with a retrograde viral vector designed to express Cre-recombinase in neurons projecting to the NAc. The DIO-Gluc vector was co-injected with a DIO-hM3Dq-mCherry vector designed to express Cre-dependent Gluc NHP-RMAs and excitatory chemogenetic receptors in BLA neurons projecting to the NAc. The injections were verified by adding GAD to the AAV suspensions and acquiring post-surgical MRI scans. Blood was collected weekly after the AAV injections to assess Cluc- and Gluc-based RMA signal levels under the hypothesis that Cluc-based signals would peak earlier than Gluc-based signals due to retrograde Cre-dependent expression for Gluc-based transgene expression. **F) Time course of Cluc- and Gluc-based RMA signal change following AAV injections**. Mean ± SEM of the Cluc- and Gluc-based RMA signal change relative to baseline across the two monkeys that received the AAV injections. The different colored asterisks above each timepoint indicate when the specific RMA signal increased above baseline (*p* < .05 Bonferroni corrected) based on a generalized linear mixed model that assessed changes in both monkeys. The inset panel depicts RMA signal change relative to baseline levels 24 hours after the AAV injections. **G) Peak RMA signal change**. Boxplots of the peak Cluc and Gluc signal change relative to baseline. Datapoints represent the peak in the timeseries of each technical replicate assessed for each RMA type and for each monkey. **H) Histological quantification of NHP-RMA markers in macaque NAc and BLA**. Cluc+ cells were quantified using in situ hybridization. Gluc+ and GFP+ cells were detected using immunofluorescence. Cell counts were normalized against DAPI+ nuclei detected within imaged regions of interest from randomly selected tissue sections spanning the anterior-to-posterior extent of the BLA and NAc, using stereotaxic injection coordinates and post-surgical MRI scans as a guide. Boxplots show the mean and distribution of cell counts within ROIs randomly sampled from each section. **I) Equivalent Cre-dependent expression of Gluc and hM3Dq-mCherry in BLA neurons**. Boxplots show the mean and distribution of the ratio of Gluc+ and mCherry+ to GFP+ cell counts within each ROI. **J and K) Representative adjacent tissue sections showing Cre-dependent gene expression of Gluc and hM3DqmCherry in BLA neurons**. Overview (4X) images showing the entire BLA and corresponding high-magnification (20X) images of Gluc or hM3Dq-mCherry (red) and GFP (green) expression in BLA neurons that project to the NAc in one monkey (A64B). * p < 0.05. For the scale bars in the J and K panels: 1 mm (4X overview) and 100 um (20X high-magnification).

We first confirmed *in vivo* Cre-dependent increases in the Gluc-based bioluminescence signal in mice. In one group of wild-type mice (*n* = 3) we co-injected AAV2-hSyn-DIO-Gluc-IRES-GFP (125 nL) and AAV5-hSyn-CRE-eGFP (125 nL). In a second group of mice we injected only the DIO-Gluc vector. Luciferase assays one and two weeks after injecting the AAVs confirmed that Cre-Lox recombination was limited to the group that was co-injected with the Cre viral vector (**Fig. 6B**; Group x Time: *F*(2,8) = 80.42, *p* < 0.0001). There was an increase in the Gluc signal from one week to the next (week 1 vs. 0: *t*(4) = 2.85, p = 0.0462; week 2 vs 1: *t*(4) = 9.28, *p* = 0.0007) in the group that was co-injected with Cre and no increase in the group that lacked Cre. Histological quantification of Gluc and GFP expression using immunofluorescence confirmed the luciferase assays (**Fig. 6C**). Gluc (Group: *t*(1) = 3.48, p = 0.025) and GFP (Group: *t*(1) = 3.45, p = 0.026) expression was only evident in the mice where the DIO-Gluc vector was co-injected with the Cre vector (**Fig. 6D**). The percentage of neurons expressing GFP was equivalent to the percentage of neurons expressing Gluc (*i*(2) = 2.15, *p* = 0.165), suggesting that co-injecting the DIO-Gluc and Cre viruses led to extensive Cre-Lox recombination.

We next evaluated in macaques if the DIO-Gluc RMA could track intersectional gene expression in basolateral amygdala (BLA) neurons projecting to the nucleus accumbens (NAc; **Fig. 6E**). There is a dense projection from BLA of the primate amygdala to NAc in the ventral striatum^23^. To express Cre-recombinase in these neurons we injected retro.AAV2-hSyn-CRE-eGFP into the ventral striatum, targeting the NAc. The retrograde viral vector was co-injected with AAV2-hSyn-Cluc (1:6 mixture) and GAD. We co-injected Cluc because retro.AAV2^24^ and AAV2 differ in their retrograde transduction^45^. We expected the Cluc vector to not undergo retrograde transport and remain in the ventral striatum. Therefore, the Cluc-based RMA signal would serve as a proxy for the earliest we could expect intersectional gene expression in BLA neurons of both Cre and DIO-Gluc.

In the same surgery we injected AAV2-hSyn-DIO-Gluc-IRES-GFP into the BLA. We co-injected the DIO-Gluc vector with AAV2-hSyn-DIO-hM3Dq-mCherry (1:1 mixture) and GAD. This allowed us to determine if the DIO-Gluc RMA signal could serve as a proxy for expression of other Cre-dependent transgenes that could be used to perturb the same targeted neural circuit. Moreover, to demonstrate the sensitivity the RMA signals in reading out gene expression once Cre was present, we injected an additional 25 ml of AAV2-hSyn-DIO-Gluc-IRES-GFP into the amygdala of each monkey 6 weeks after the initial AAV injections.

We hypothesized we would observe early Cluc-based increases in RMA signal since there were no constraints on Cluc transgene expression and delayed Gluc-based increases in RMA signal due to constraints imposed by retrograde Crelox recombination.

Consistent with our predictions, increases in the Cluc and Gluc RMA signals differed in time (Serotype x Time: *F*(11, 153) = 11.22, *p* < 0.0001; **Fig. 6F and G**). As we described earlier, the Cluc signal increased 24 hours after the injection compared to baseline (*p* = 0.0121 corrected) and was followed by a decrease in the Cluc signal at one week. The Cluc signal then increased relative to baseline from two until eight weeks (all *p* < 0.01 Bonferroni corrected) after the injection and peaked with a 1.45-fold increase at 4 weeks before the signal started to decline. The DIO-Gluc signal after 24 hours was not increased relative to baseline (*p* = 0.22 corrected). Rather there was a moderate 1.16-fold increase in the DIO-Gluc signal relative to baseline one week after the initial injection surgery that slowly increased over time (Time: *F*(11,62) = 13.82, *p* < 0.0001), with a peak 1.41-fold increase at 9 weeks (**Fig. 6F**).

For one of the two monkeys that we injected with the DIO-Gluc vector we randomly selected tissue sections from the monkey’s brain surrounding the stereotaxic coordinates where we injected each of the AAVs and specified target and control regions of interest (ROI). Using in situ hybridization to detect Cluc (*t*(9) = 6.50, *p* = 0.0001) we confirmed expression in ventral striatal neurons within the injected (*M* = 1.74%) compared to control ROIs (**Fig. 6H** and second row (A64B) in **Fig. 5E**). Using immunofluorescence we confirmed heightened expression of Gluc (*M* = 1.43%; *t*(6) = 2.43, *p* = 0.0513), GFP (*M* = 3.35; *t*(12) =5.21, *p* = 0.0002), and mCherry (*t*(5) = 3.23, *p* = 0.0231) in BLA neurons (**Fig. 6H**). Because we used a commercially available retrograde Cre-eGFP vector to transfect amygdala neurons projecting to striatum we couldn’t readily disambiguate GFP expression driven by the Cre vector compared to the DIO-Gluc vector. We did detect fewer Gluc+ cells compared to GFP+ cells (*t*(13) = −2.16, *p* = 0.049). We could, however, disambiguate Cre-eGFP expression from mCherry expression linked to expression of hM3Dq chemogenetic receptors in amygdala neurons. We similarly found fewer mCherry+ than GFP+ neurons (t(17) = −2.34, *p* = 0.0318). We computed the ratio of Gluc+ and mCherry+ neurons to GFP+ neurons (**Fig. 6I**) in the same region of interest. We found statistically equivalent expression of Gluc+ and mCherry+ neurons (*F*(1,11) = 0.356, *p* = 0.563) consistent with us injecting similar quantities of the DIO-Gluc and DIO-hM3Dq-mCherry vectors into the amygdala. Similar expression of these two unrelated Cre-dependent transgenes injected in similar quantities implies that the Gluc-based signal can function as a readout of Cre-lox recombination. This also validates retrograde expression of Cre via the retro.AAV2 capsid^45^. The DIO-Gluc vector can be co-injected with other viral vectors and used as a proxy for confirming expression of functional transgenes that enable neuronal and circuit perturbations. It is encouraging that the Gluc-based signal quickly doubled after a second injection of the DIO-Gluc vector into the amygdala after 6 weeks of expression. However, when we co-injected DIO-Gluc and Cre vectors in mice and in macaques, macaques showed relatively lower Gluc signals in the blood and lower Gluc+ neuronal expression. This underlines the difficulty in using intersectional viral tools to express transgenes in spatially distributed projection neurons in the larger primate brain.

## DISCUSSION

The ability to repeatedly monitor transgene expression in the nonhuman primate brain is critical for the continued development, validation, and use of molecular and genetic tools in NHPs. Here, we show that a simple blood draw can monitor expression of transgenes in the NHP brain. To achieve this, we engineered a new reporter based on a recent technology called Released Markers of Activity, or RMAs^10^. While RMAs were originally designed to cross from the brain into the blood of mice, we found that a simple modification of these markers was sufficient to enable similar reverse transcytosis and transgene expression monitoring in the brains of NHPs. To accommodate species differences, we modified the mouse Fc domain in the RMA to its equivalent in rhesus macaque. We found that the purified NHP-RMA proteins could enter the blood at detectable levels within minutes after an intracranial injection. We also found detectable levels of NHP-RMAs in the blood when the NHP-RMA protein was secreted by neurons in one or more brain regions following an intracranial injection of designed AAVs. This occurred as early as 24 hours after the start of the injection and continued up until 8-12 weeks after the initial gene delivery. By expressing the floxed RMA construct in BLA neurons projecting to the NAc that co-expressed the Cre enzyme, we found that NHP-RMAs monitor the progression of Cre-lox recombination-dependent transgene expression in a specific neural circuit. We detected a gradual, specific increase of the Cre-dependent Gluc-based RMA signal in the blood over the course of weeks. Finally, we show that NHP-RMA detection can be multiplexed. By designing two variants of NHP tailored RMAs with different luciferases, we could monitor transgene expression simultaneously from either two brain regions or two distinct neuron types in the same animal, as previously demonstrated in rodents. Importantly, the NHP-RMA serum markers were detectable when using only 5 microliters of serum and a standard microplate luminometer that is available in most labs and research centers.

The ability to repeatedly monitor transgene expression using RMAs via simple blood draws has several advantages. First, it enables researchers to be confident that the viral vectors injected into the brain are functioning as intended. At present, NHP researchers have few good options to confirm success of gene delivery *in vivo* in the studied animal. In some cases, it is possible to use expensive neuroimaging approaches (e.g. rs-fMRI or PET) to visualize expression of some transgenes, which require development and use of contrast agents for these modalities^7,25–27^. Repeated monitoring also requires repeated anesthetic events. More commonly, however, the analysis is done post-hoc after animals are euthanized through histology. In the case of such post-mortem analysis the researchers have little to no information about the temporal progression of expression and how that progression would affect the results. By co-expressing RMAs with the delivered genetic cargo, researchers can use a simple blood draw to assess the success and temporal progression of transduction, enabling correlations between expression levels and behavioral or electrophysiological outcomes. For example, nonhuman primate neuroscientists could use RMAs to validate experimental manipulations without the need for euthanasia. RMAs have the potential to refine transgene monitoring practices—saving time and resources—while most importantly maximizing the scientific contributions of individual NHPs. By enabling the planning of longer-term or hierarchical experiments, RMAs can increase the overall utility of nonhuman primates in both clinical and systems neuroscience research.

An interesting discovery that will require further research is the rapid expression of the markers. We found detectable RMA signal at 24 hours after AAV delivery, despite the commonly reported gradual AAV expression of genetic cargo over several weeks^28^. Further work will be needed to understand the mechanisms underlying this rapid expression. One possibility is that the second strand synthesis^29^ is not needed for this initial bout of expression. Positive and negative strands of multiple AAVs pairing to create functional dsDNA, before being separated and routed towards second strand synthesis. *In vitro*, dsDNA can be created within 3 hours following transfection^22^, suggesting an early bout of expression is possible.

The maximum RMA signal in the blood was achieved within 4 to 6 weeks. NHP systems neuroscientists typically begin experiments requiring AAV-mediated transgene expression 8 to 12 weeks after AAV delivery ^30–32^. Our results suggest a shorter timeline could be and may even be beneficial as we observed drops in the overall RMA expression from 8 to 12 weeks after initial AAV delivery. All the AAVs we injected in the NHPs involved the AAV2 capsid. One future application of RMAs is high throughput screening of different AAV serotypes to determine if they modulate the time course of transgene expression.

One potential limitation of the NHP-RMA is its level of maximum signal. While the signal was detectable, the signal over baseline levels were 1-2 orders of magnitude smaller than those observed in mice. One possible reason could be less effective transport of NHP-RMA to the macaque’s neonatal Fc receptor (FcRN) that mediates reverse transcytosis across the BBB. For example, in nonhuman primates the serum bioluminesence signals in the blood following injections of AAVs encoding RMAs is slower to reach asymptote, suggesting fewer proteins enter the blood per second than in mice. This suggests that the Fc region of the macaque IgG1 may need to be modified, or Fc fragments from other antibodies may need to be tested to improve the reverse transcytosis. There are multiple IgG Fc receptor classes (**Fig. 2A**) in the nonhuman primate brain and functional homologies between the rodent and human Fc receptor classes are not well established. Future experiments evaluating the relative efficacy of RMAs targeting different nonhuman primate Fc receptors should aid in optimizing RMA sensitivity, signal level, and pharmacokinetics. This is an important challenge as a higher signal-to-noise ratio will likely be required for using RMAs to monitor small changes in gene expression, such as using NHP-RMA variants to measure c-Fos levels in NHP brains, spying on rare cell populations, or transcript levels of other genes through RNA sensors^12-14,33^.

A second limitation of the current RMA variants is the eventual loss of the blood-derived bioluminesence signals over time. We observed a progressive increase in RMA detection over the first 8 weeks following intracranial viral injections, however, after 8-10 weeks we saw a decline in bioluminesence signals back towards or equivalent with baseline measurements. We can only speculate about why this occurs. When we injected the purified NHP-RMA protein directly into the brain we found a strong, positive dose-dependent correlation with the amount injected and the RMA signal in the blood (**Fig. 3D**). One possibility is that buildup of the RMAs in the blood over weeks triggers an immunogenic response leading to the production of neutralizing antibodies against the RMAs. This may occur similarly to how antibody based therapeutics can lose potency due to anti-drug-antibodies^34^. Prior studies examining the use of RMAs in rodents have only evaluated their expression out to 5 weeks^16^. Identifying the mechanism underlying this drop off in RMA signals is another important challenge to using RMAs for long-term monitoring of transgene expression. The ability to verify initial transgene expression over the course of eight weeks should still facilitate increased use of viral tools in systems and clinical neuroscience studies involving NHPs.

Future development of the RMA paradigm and its application to NHPs will likely involve multiple improvements and applications. For example, recent work shows that RMAs can be tied to expression of RNA-based sensors^12,13,33^ to monitor transcript level of theoretically any chosen gene^14^. The same work recorded expression of immediate early genes such as c-Fos or Arc^14^ as proxies of neuronal activation. Improving the sensitivity of NHP-RMAs will be crucial to observe gene expression dynamics and take advantage of ever-increasing information about the genetic diversity of neuron types in the primate brain^35,36^. A helpful innovation would be to temporarily reduce or erase RMA signal levels in the blood just before blood is drawn to enable a baseline reset. This would improve the dynamic range of RMA measurements in many contexts and was recently demonstrated in rodents through the use of protease-digestible RMAs (fast-erasable RMAs, feRMAs)^37^.

Since RMAs are proteins, and their detection can be done by any relevant biochemical method. Massively multiplexed detection of barcoded RMAs is possible for example through use of mass-spectrometry^38^, single-molecule protein sequencing^39,40^, or even antibody-based assays such as olink^41^. Improving the multiplexing of RMA detection could allow one to study multiple brain regions of a single animal, or expression of multiple genes. The potential for high multiplexity is an important advantage of RMAs design over optical imaging, MRI, or PET. These neuroimaging techniques can only record up to several signals at once, compared to thousands of protein barcodes that can be detected with mass spectrometry or other proteomic assays.

Finally, extending the duration of expression of RMAs could be valuable for long-term studies in slow developing disease models of cancer, neurodegenerative conditions, or aging. One potential improvement would be to use syngeneic proteins as RMAs, or other proteins. Any protein that can be secreted from neurons, cross the BBB, and be easily detected in the blood can act as an RMA. Thus, multiple types of reporters could be tested beyond luciferases from *Gaussia* or *Cypridina*, as presented in this study. Another option could be to avoid constant exposure of the system to RMAs and thus reduce potential immunization, by providing inducible protein expression that can be induced occasionally to a minimum detectable level, whenever measurement is needed. In this study RMAs were expressed continuously, even when the blood was not drawn.

RMAs are primarily intended as a research tool. However, they may also be useful in the clinic. Gene therapy or cell transplantation often lack tools needed to evaluate the duration or success of transduction in deep tissues. By co-dosing low, but detectable, levels of RMAs, one could monitor the success and duration of these therapies as proxies. Likewise, in psychiatry the impact of neuromodulatory treatments (e.g. TMS or tFUS) on specific cell types or neural circuits that are beyond the resolution of neuroimaging assessments can be repeatedly, accessibly, and frequently monitored with a simple blood test. With such an approach, it would be possible to better evaluate the reason for failure of these therapies or separate responding from non-responding clinical trial participants.

Overall, these results demonstrate the feasibility of using engineered serum markers for monitoring transgene expression in the nonhuman primate brain and substantiate further optimization of these tools with a future goal of non-invasively monitoring endogenous gene expression in the primate brain as it relates to changes in structure and function.

## MATERIALS AND METHODS

### Animal subjects

#### Mice

Wild-type C57BL/5J (strain 000664) male and female mice were purchased from The Jackson Laboratory and used for experiments at 10-12 weeks of age. Mice were housed with a 12 hr light/dark cycle, 18-23°C ambient temperature, and 40-60% humidity and were provided with food and water ad libitum. Mice experiments were performed under the protocol approved by the Institutional Animal Care and Use Committee of Rice University.

#### Rhesus macaques

Adult rhesus macaques (*n* = 9; 6 female) of Indian origin were acquired and housed at the Oregon National Primate Research Center (ages 4.1-12.2; 5.9-13.1 kg; Table S1). Animals were pair-housed when possible or otherwise singly housed and maintained on a 12 hr light-dark cycle. All animals had ad libitum access to water and ad libitum access to chow. The Oregon National Primate Research Center and Oregon Health and Science University Institutional Animal Care and Use Committee and Institutional Biosafety Committee approved all procedures and protocols and all of the guidelines specified in the National Institutes of Health Guide for the Care and Use of Laboratory Animals were followed.

#### Plasmid construction

Protein sequence of Rhesus macaque IgG1 heavy chain region was obtained from Protein Data Bank (PDB) ID: 6D4E. The sequence showed identical for both Macaca mulatta (NCBI accession No. ATV90900.1) and Macaca fascicularis (NCBI accession No. AFF60181.1). To construct pET-T7-His-Gluc-NHP.RMA, the DNA sequence was codon-optimized for *E. coli* expression and synthesized as a gBlock gene fragment in Integrated DNA Technologies. The DNA segment encoding Gluc was amplified by PCR using our previous mouse-specific construct pAAV-hSyn-Gluc-RMA-IRES-EGFP (Addgene #189629) as a template and was purified using the Monarch DNA Gel Extraction Kit (New England Biolabs). The two inserts, gBlock encoding the macaque IgG1 Fc region and Gluc segment, were inserted into a linearized pET28a vector using Gibson Assembly.

To construct pAAV-hSyn-Gluc-NHP.RMA-IRES-eGFP, the same vector pAAV-hSyn-Gluc-RMA-IRES-EGFP (Addgene #189629) was digested with KpnI and EcoRV (New England Biolabs) to isolate the linearized backbone. Two inserts, Gluc and IRES-eGFP, were amplified from the same vector and the macaque IgG1 Fc insert was amplified from the synthesized gBlock. All three inserts were then assembled with the backbone using Gibson Assembly. Using a similar strategy, pAAV-hSyn-Cluc-NHP.RMA was constructed by digesting pAAV-hSyn-Cluc-RMA (Addgene #189624) to obtain the vector backbone and inserting the amplified Cluc and macaque IgG1 Fc segments through Gibson Assembly. To construct hSyn-DIO-Gluc-NHP.RMA-IRES-eGFP, pAAV-hSyn-DIO-Gluc-RMA-IRES-EGFP (Addgene #189630) was digested with NcoI and its vector backbone was isolated through gel extraction. DNA segment encoding Gluc-NHP.RMA-IRES was amplified from the construct pAAV-hSyn-Gluc-NHP.RMA-IRES-eGFP made in this study and inserted in inverted orientation into the backbone to ensure Cre-dependent expression.

#### Protein purification

Purification was performed similarly to our previous study with slight modifications^10^. Shuffle T7 Express chemically competent *E. coli* cells (New England Biolabs #C3029J) were transformed with plasmid pET-T7-His-Gluc-NHP.RMA. The next day, a single colony was picked and grown overnight in 3 ml of LB medium at 30°C with shaking at 250 RPM. The culture was then transferred to 1 L of Terrific broth and incubated under the same condition until the OD600 reached 0.5. The culture was cooled on ice for 30 min, and 100 µM IPTG was added to induce expression for 20 hrs at 16°C with shaking at 180 RPM. Cells were pelleted by centrifugation, resuspended in lysis buffer (300 mM NaCl, 50 mM NaH2PO4, 10 mM imidazole, 10% glycerol, pH 8.0) supplemented with ProBlock Gold protease inhibitor (Gold Biotechnology), and lysed by a sonicator (VCX 130, Sonics and Materials). Lysates were centrifuged at 12,000 x *g* for 30 min at 4°C and the resulting supernatant was subjected to binding with Ni-NTA agarose resins (Qiagen #30210) and loaded into chromatography columns (Bio-Rad). The resins were washed by gravity flow using lysis buffer containing increasing concentrations of imidazole up to 50 mM. Proteins were eluted with lysis buffer containing 500 mM imidazole and buffer-exchanged into PBS using an Amicon centrifugal filter unit with a 30 kDa cutoff (MilliporeSigma). Proteins in PBS were analyzed by SDS-PAGE. To increase purity, NHP-RMA proteins were isolated using a HiLoad Superdex 75 pg size exclusion chromatography column (Cytiva #28989333) connected to an AKTA Pure25 FPLC (Cytiva), and the peak fractions were pooled and further concentrated in PBS. A BCA protein assay (Thermo Fisher Scientific #23225) was used to determine protein concentration.

#### Adeno-associated virus production

Constitutive expression plasmids encoding Gluc-NHP.RMA and Cluc-NHP.RMA, along with a Cre-dependent expression plasmid for Gluc-NHP.RMA, were amplified using a PureYield Plasmid Maxiprep kit (Promega #A2393). SmaI digestion was performed to verify the integrity of the inverted terminal repeats of the amplified plasmids. AAVs with serotype 2 were produced for all the above constructs by the BRAIN Initiative Neurotools Viral Vector Core located at Duke University. AAV titers were obtained as follows: 7.87×10^12^ vg ml^−1^ for AAV2-hSyn-Gluc-NHP.RMA-IRES-eGFP; 1.62×10^13^ vg ml^−1^ for AAV2-hSyn-Cluc-NHP.RMA; and 2.17×10^13^ vg ml^−1^ for AAV2-hSyn-DIO-Gluc-NHP.RMA-IRES-eGFP. AAV5 and retro.AAV2 encoding pENN.AAV.hSyn.HI.eGFPCre.WPRE.SV40 was obtained from Addgene (Addgene #105540-AAV5 and #105540-AAVrg) at respective titers of 1.76×10^13^ vg ml^−1^ and 1.7×10^13^ vg ml^−1^ encoding. All AAVs were stored at –80°C after aliquoting into smaller quantities and kept on wet ice during surgery until they were injected.

#### PC-12 transduction

PC-12 (ATCC #CRL-1721) was cultured in RPMI 1640 medium (Corning #10-040-CV) supplemented with 10% horse serum (Life Technologies #26-050-088) and 5% fetal bovine serum (FBS) (Corning #35-011-CV). Cells were incubated in humidified air with 5% CO_2_ at 37°C and split every 2 d with a subcultivation ratio of 1:2 or 1:3. To transduce PC-12, a 96-well plate was coated with 10 µg ml^−1^ of collagen IV (Corning #354233) in PBS at 4°C overnight, then dried under UV light. PC-12 was seeded at 10,000 cells per well in the coated plate. After 20 hrs, AAVs were added into each well to transduce the cells at a multiplicity of infection at 100,000 (1 × 10^9^ vg per well). For example, AAV2-hSyn-Gluc-NHP.RMA-IRES-eGFP was diluted tenfold (7.87 ’ 10^11^ vg ml^−1^) and 1.3 µL was added to the well. After 5 d, the supernatant was collected after centrifuging the media samples at 500 x *g* for 3 min to exclude the cell pellets and stored in –20°C until use.

### Luciferase assay

Gluc substrate, 0.5 mM native coelenterazine (CTZ) stock (Nanolight Technology #303) was dissolved in luciferase assay buffer (10 mM Tris, 1 mM EDTA, 1.2 M NaCl, pH=8.0) containing 66% DMSO and stored at –80°C. For Cluc substrate stock, 50 µM of vargulin (Nanolight Technology #305) was dissolved in acidic n-butanol and stored at –80°C. Before measuring bioluminescence, the CTZ and vargulin stocks were diluted to 20 µM and 1.0 µM, respectively, in luciferase assay buffer and kept in dark at room temperature for 1 hr. Media samples were thawed and 25 µL was transferred to a black 96-well plate (Corning). Infinite M Plex microplate reader equipped with icontrol software (Tecan) was used to inject 50 µl of the assay buffer containing CTZ or vargulin and to measure photon emission integrated over 30 s. The values were averaged to calculate the light unit per second.

### Mouse stereotaxic intracranial viral injections

Mice were anesthetized in 1.5%-2% isoflurane in air or O2. The scalp was disinfected with chlorohexidine scrub and solution before a longitudinal incision was made. AAVs were injected into the mice brains using a 1 microliter syringe equipped with a 34-gauge beveled needle (Hamilton) attached to a motorized pump (World Precision Instruments) and a stereotaxic frame (Kopf). 125 nl of each AAV was injected with the following viral doses (using the estimated stock titers): 9.84×10^8^ vg of AAV2-hSyn-Gluc-NHP.RMA-IRES-eGFP; 2.03×10^9^ of AAV2-hSyn-Cluc-NHP.RMA; 2.71×10^9^ vg of AAV2-hSyn-DIO-Gluc-NHP.RMA-IRES-eGFP; and 2.20×10^9^ vg of AAV5-hSyn.HI.eGFP-Cre.WPRE.SV40. A total volume of 250 nl of AAVs, containing 125 nl of each AAV, was unilaterally injected into the Cd/Pu of the left hemisphere (AP +0.25 mm, ML +2.0 mm, DV –3.2 mm) infused at a rate of 250 nl min^−1^, and the needle was kept in place for 5 min before taking it out from the injection site. Following the injection, the incision was closed with GLUture topical tissue adhesive (World Precision Instruments) and the mice were monitored for at least 5 d post-operatively.

### Macaque stereotaxic intracranial injections of purified NHP-RMA protein and AAVs Neuroimaging

On the day of surgery, all monkeys received MRI scans to determine surgical coordinates for the AAV injections. Anesthesia was first induced with ketamine (10 mg/kg IM), followed by maintenance anesthesia with isoflurane gas vaporized in 100% oxygen. Animals were placed in a MRI-compatible, NHP-specific stereotaxic frame (Crist) in which they remained for the duration of the scan and subsequent surgery. A T1-weighted scan was collected on a Siemens Prisma scanner using a surface coil. Brain images were subsequently examined using AFNI^42^. The injection coordinates were selected and transferred from MRI-space to stereotaxic surgical space. Monkeys were taken directly from the MRI to the operating room following their scan and maintained under anesthesia. If collected, post-operative MRI scans followed the same procedure to verify successful injection of the AAVs and assess targeting.

### Stereotaxic Surgery

Pre-operative care consisted of overnight food restriction for 12 hrs and a general examination to ensure that patient health was adequate to withstand the procedure. All subsequent described procedures were performed under aseptic conditions. Local anesthetic was injected subcutaneously along the incision site and the scalp was prepared for incision. Depending on the number and size of the targeted brain regions, between one and eight small craniotomies (each ~0.5 cm in diameter) were made using an air drill to expose the dura (**see Table S1 and S2**). The dura was incised to allow the needle to penetrate into the brain. A 100 µl Hamilton syringe fitted with a 27-gauge needle (blunt tip) was used for injections into the basal ganglia and cortex. A 30-gauge sheathed needle was used for injections into the amygdala (**see Table S1 and S2**). The Hamilton syringe was connected to a programmable stereotaxic injection pump (Harvard Apparatus) mounted to the micromanipulator of the stereotaxic arm. The syringe was first primed by twice loading and expelling 80-100 µl of the injectate (either the purified protein or AAV suspension). The injectate used to prime needle was then either mixed into the final injectate or discarded. The purified NHP-RMA protein or an AAV suspension was injected through the syringe. The infusion rate was programmed to begin at either 0.2 ul/min (cortical injections) or 0.5 µl/min (basal ganglia and amygdala injections) and ramped linearly to 1.5 uL/min (cortical injections), 2.5 uL/min (amygdala injections), or 3.5 µl/min (basal ganglia injections) depending on the injection site target and total volume of the injection (**see Tables S1 and S2**). This progressive increase in infusion rate is known as convection enhanced delivery, and it improves the spread of infusate compared to conventional infusion methods. Depending on the brain region injected, multiple injections were made along the dorsoventral axis of the needle track starting from the most ventral target. After each injection, the needle was left in place for an additional 5 min to allow the injectate to diffuse from the needle tip before removing^43,44^.

After microinjections were completed, the skull opening was filled with gelfoam, the incision was closed, and the animal was monitored closely during recovery. Post-operatively, animals were monitored for 5–7 days by veterinary staff and received Cefazolin, Hydromorphone, and Buprenorphine (antibiotic and pain management). All surgical procedures were conducted by trained Surgical Services Unit personnel or trained laboratory staff under the supervision of surgical veterinarians in dedicated surgical facilities using aseptic techniques and comprehensive physiological monitoring.

Intracranial injections of the purified NHP-RMA protein were performed in four monkeys (all female) and targeted the posterior putamen (**see Table S2**) and intracranial viral injections of the AAVs encoding the NHP-RMAs were performed in five monkeys (1 male; **see Table S3**). Unless otherwise noted otherwise all AAVs were diluted to achieve a titer of 1×10^12^ vg/mL.

### Blood collection for luciferase assay

#### Mice

Mice were anesthetized in 1.5%-2% isoflurane in air or O2. Following, 1-2 drops of 0.5% ophthalmic proparacaine were applied topically to the cornea of an eye. Heparin-coated microhematocrit capillary tube (Fisher Scientific #22-362566) was placed into the medial canthus of the eye and the retroorbital plexus was punctured to withdraw 50-100 µl of blood. The collected blood was centrifuged at 1,500 x *g* for 5 min to isolate serum. Samples were stored at −80°C until analysis.

### Macaques

Intraoperatively, whole blood was collected from an IV catheter. Prior to collection, the IV catheter was flushed with sterile saline to ensure patency, 3 mL of waste was aspirated from the IV catheter, then a sample of 1.5 mL of whole blood was collected. Post-operatively, whole blood was collected via venipuncture following standard blood draw procedures in either a bleed tower or following a standard ketamine anesthesia protocol. Samples were collected into a 2 mL clear microcentrifuge tube with no additive or anticoagulant. Following collection, samples were immediately centrifuged for 5 m at 22°C and 3,000 RPM. Blood serum was removed using a 1000 uL pipette and transferred into a 1.5 mL black microcentrifuge tube and flash frozen on dry ice. Samples were stored at −80°C until analysis.

### Luciferase assay

In each species 5 µl of serum was mixed with 45 µl of PBS + 0.001% Tween-20 in a black 96-well plate to conduct a luciferase assay. Bioluminescence was measured using the microplate reader by injecting 50 µl of 20 µM CTZ or 1.0 µM vargulin dissolved in the luciferase assay buffer to measure Gluc or Cluc activity, respectively.

### Half-life measurement

Two monkeys received IV injections of purified NHP-RMA proteins. One monkey received 0.33 mg/kg (high dose) and the other 0.02 mg/kg (low dose). Blood samples were collected at multiple time points (5, 10, 15, 40, and 60 minutes; 24 hrs; daily or weekly out to two weeks) and processed as described above. To determine the pharmacokinetic parameters, halflives of the distribution (*α*) and elimination (*β*) phases in the two-phase clearance model were calculated by applying loglinear regression to the measured bioluminescence signals. Time points before and after 24 hrs were used to calculate *a*- and *b*-phase half-lives, respectively.

### In situ hybridization

Probes targeting *Cypridina* luciferase mRNA were designed and obtained from Molecular Instruments. The in situ hybridization chain reaction (HCR) was performed according to the manufacturer’s protocol (Molecular Instruments). Briefly, mice and macaque brain sections were incubated overnight at 37°C with binding probes in the supplied hybridization buffer. After washing with the probe wash buffer, sections were incubated overnight at room temperature with amplifier hairpins in the amplification buffer. Finally, mice and macaque sections were mounted on glass slides using mounting medium (Vector Laboratories) and imaged using a fluorescence microscope (Keyence).

### Histological imaging and analysis

#### Mice

Mice brains were extracted and postfixed in 10% neutral buffered formalin (Sigma-Aldrich #HT501128). Coronal sections were cut at a thickness of 50 µm using a vibratome (Leica). Sections were stained as follows: 1) block for 2 hrs at room temperature with blocking buffer (0.2% Triton X-100 and 10% normal donkey serum in PBS); 2) incubate with primary antibody overnight at 4°C; 3) wash in PBS for 15 min 3 times; and 4) incubate with secondary antibody for 4 hrs at room temperature. After last washes in PBS, sections were mounted on glass slides using the mounting medium (Vector Laboratories). Antibodies and dilutions used are as follows: rabbit anti-Gluc (Nanolight Technology #401P): 1:1000; donkey ant-rabbit Alexa Fluor 594 secondary antibody (Life Technologies): 1:500. Mice images were acquired with a BZ-X800 fluorescence microscope (Keyence). Manual cell counting was performed using the Zen Blue software (Zeiss) by blinded observers uninformed of the experimental conditions. The DAPI positive cells in the images taken at CP were quantified by dividing the area into 16 subregions (4 by 4) using gridlines, randomly selecting 4 subregions for cell counts, and multiplying by 4 to estimate the total number of positive cells. Cells in all other images were counted individually.

#### Macaque

Necropsies and tissue collections were performed as previously described^45,46^. Animals were sedated with ketamine (10 mg/kg) and then deeply anesthetized with sodium pentobarbital followed by exsanguination. The brain and spinal cord were perfused through the ascending carotid artery with 1 L of 0.9% ice cold saline. The brain was then removed from the skull (< 30 min post-mortem), deposited into an icecold bath of saline for transport, and placed into an ice-cold, steel brain matrix (Electron Microscopy Sciences). A carbon steel knife blade (Thomas Scientific) was inserted into the first slot of the brain matrix and additional blades inserted in 4 mm increments. Depending on the size of the brain this resulted in 18 to 24 brain slabs that encompassed the anterior-posterior extent of the macaque brain. The resulting bilateral brain slabs were then removed from the brain matrix and drop fixed in 4% PFA (prepared from 32% concentrate diluted in 0.1M phosphate buffer) for 48 hrs. The fixed tissue was then cryoprotected in 10% sucrose with the concentration increasing up to 30% sucrose over 3-5 days. All tissue was stored in 30% sucrose long term or until analysis.

Cryoprotected tissue was sliced at 40 uM on a freezing microtome. Native autofluorescence was quenched under a 6,000 lumen LED light at 4°C in 1:1 solution of 0.05M PBS and diH2O for 48 hrs^47^. Tissue sections were rinsed in a solution of Tris-buffered saline (TBS) and 0.05% Triton X-100 (DM) and then blocked in 5% normal goat serum (Gibco). Sections were incubated in a primary antibody solution overnight at room temperature or for 48 hrs at 4°C using primary antibodies against Gluc (Prolume Ltd.; mouse anti-Gluc: 1:500; mouse anti-Gluc: 1:300), GFP (Aves Labs, Inc.; chicken anti-GFP: 1:500), nuclei using DAPI (Hello Bio; 1:1000 in PBS), and mCherry (AbCam; rabbit anti-mCherry: 1:500). Sections were sequentially incubated in secondary antibody solutions for 1 hr at room temperature. After each incubation the sections were washed in DM prior to beginning the next incubation: 1) Sections were incubated in goat anti-chicken Alexa Fluor 488 (Invitrogen; 1:500) conjugated secondary antibody solution; 2) Sections were incubated in a goat secondary antibody (anti-rabbit: 1:500; anti mouse: 1:300) conjugated with a fluorophore (either Alexa Fluor 568 or Alexa Fluor 647); 3) Sections were incubated in a DAPI solution for 10 min at room temperature before subsequent washes in TBS. Processed slices were mounted onto microscope slides using ProLong™ Diamond Antifade Mountant (Invitrogen).

Images were taken on a Keyence BZ-X710 microscope. 4X images were stitched together using the BZ-X Analyzer to provide slice overviews. 20X z-stack images were acquired, ranging from 10-35 uM in thickness with a pitch of 0.4 uM. The BZ-X Analyzer was used to generate an extended depth of field (EDF) image and optimize gain and exposure settings. All DAPI positive cells in each 20X image were detected using an automated algorithm in QuPath^48^. Cells expressing other protein markers were detected visually and the counted by two different experimenters whose counts were averaged.

#### Fc sequence alignment

Rhesus macaque Fc sequences of IgG1, IgG2, IgG3, and IgG4 were obtained from Protein Data Bank (PDB) with ID numbers 6D4E, 6D4I, 6D4M, and 6D4N, respectively. Mouse IgG1 Fc sequence was also obtained from our previous study of mouse-specific Gluc-RMA. The five Fc sequences were aligned using the Jalview 2.11.3.3 software. Conservation score for each residue position was calculated by the software based on the similarity of physicochemical properties (charge; polarity; size; hydrophilic vs. hydrophobic; and aromatic vs. aliphatic), with a score of 10 indicating complete conservation of all properties^49^.

### Statistical analysis

All statistical comparisons were performed using JMP Pro (17.0.0; SAS) and MATLAB (2024b; Mathworks).

### PC-12 tissue culture

Time dependent increases in RMA signal levels in the tissue culture media for each of the NHP-RMA AAVs were compared across baseline and post-transfection tissue samples using a generalized liner mixed model that specified each independent tissue culture as a random effect and the baseline and post-transfection timepoints as a fixed effect. For the model comparing the effects of transfecting the tissue culture with AAV encoding DIO-Gluc, each independent tissue culture was included as random effect. Fixed effects included the presence or absence of Cre baseline vs. post-transfection timepoints. Full models including all relevant interactions between random and fixed effects were used. Post-hoc comparisons used Tukey’s correction for multiple comparisons when more than three post-hoc comparisons were performed.

### Mouse and macaque luciferase and histological assays

Time dependent increases in RMA signal levels in mice and macaques after AAV or protein injections were compared across baseline and post-injection serum samples using a generalized liner mixed model that specified each technical replicate within each animal as a random effect and timepoints as a fixed effect. Group effects pertaining to Cre expression and RMA serotype were also modeled as fixed effects when appropriate. Histological analyses of Gluc, Cluc, and GFP expression in both species specified individual target and control region of interests located within individual tissue sections as random effects nested with the individual tissue section and animal identity. The region of interest, hemisphere, or marker protein identity were specified as fixed effects based on the experimental design. Post-hoc comparisons used Tukey’s or Bonferroni corrections for multiple comparisons when more than three post-hoc comparisons were performed.

## Data Availability

The authors declare that all data supporting the results of this study are available within the paper and its Supplementary Information. Raw and analyzed datasets are available from the corresponding author upon reasonable request. The plasmids designed in this study will be deposited and publicly available in Addgene or available through the Neurotools Viral Vector Core.

## ACKNOWLEDGMENTS

This research was supported by the David and Lucile Packard Foundation 2021-73005 (JOS.), and the National Institutes of Health awards R01 MH125824 (VDC), P51 OD011132 (VDC), P51 OD011092 (VDC). We thank members of the Division of Comparative Medicine, Surgical Services Unit, and Pathology Services Unit at ONPRC for assistance with intracranial injections, blood sample collection, and tissue collection. Schematic illustrations were generated using Biorender, Adobe Illustrator, and Affinity Designer.

## AUTHOR CONTRIBUTIONS

Conceptualization: VDC, JOS

Methodology: SL, MDR, VDC, JOS

Formal analysis: MST, ES, VDC, RCJ

Investigation: SL, MDR, SH, MC, JOS, VDC

Resources: JOS, VDC

Data curation: SL, MDR, SW, JOS, VDC

Writing-original draft: SL, JOS, VDC

Writing-review and editing: SL, MDR, JOS, VDC

Visualization: SL, SW, MDR, VDC, ER, HL

Supervision: JOS, VDC

Project administration: SL, MDR, JOS, VDC

Funding acquisition: JOS, VDC

