## Supplemental Materials for "Synthetic Serum Markers Enable Noninvasive Monitoring of Gene Expression in Primate Brains"

**Supplemental Information and Data**

**SUPPLMENTAL INFORMATION**

**Information S1. NHP.RMA protein sequences.**

**Gluc-NHP.RMA**

MGVKVLFALICIAVAEAKPTENNEDFNIVAVASNFATTDLDADRGKLPGEKLPLEVLKELEANARKAGCTRGCLIC

LSHIKCTPKMKKFIPGRCHTYEGDKESAQGGIGEAIDDIPEIPGFKDLEPIEQFIAQVDLCVDCTTGCLKGLANVQ

CSDLLKKWLPQRCATFASKIQGQVDKIKGAGDDGSLLGGPSVFLFPPKPKDTLMISRTPEVTCVVVDVSQEDP

DVKFNWYVNGAEVHHAQTKPRETQYNSTYRVVSVLTVTHQDWLNGKEYTCKVSNKALPAPIQKTISKDKGQP

REPQVYTLPPSREELTKNQVSLTCLVKGFYPSDIVVEWESSGQPENTYKTTPPVLDSDGSYFLYSKLTVDKSRW

QQGNVFSCSVMHEALHNHYTQKSLSVSPGK

**Cluc-NHP.RMA**

MKTLILAVALVYCATVHCQDCPYEPDPPNTVPTSCEAKEGECIDSSCGTCTRDILSDGLCENKPGKTCCRMCQ

YVIECRVEAAGWFRTFYGKRFQFQEPGTYVLGQGTKGGDWKVSITLENLDGTKGAVLTKTRLEVAGDIIDIAQA

TENPITVNGGADPIIANPYTIGEVTIAVVEMPGFNITVIEFFKLIVIDILGGRSVRIAPDTANKGMISGLCGDLKMME

DTDFTSDPEQLAIQPKINQEFDGCPLYGNPDDVAYCKGLLEPYKDSCRNPINFYYYTISCAFARCMGGDERASH

VLLDYRETCAAPETRGTCVLSGHTFYDTFDKARYQFQGPCKEILMAADCFWNTWDVKVSHRNVDSYTEVEKV

RIRKQSTVVELIVDGKQILVGGEAVSVPYSSQNTSIYWQDGDILTTAILPEALVVKFNFKQLLVVHIRDPFDGKTC

GICGNYNQDFSDDSFDAEGACDLTPNPPGCTEEQKPEAERLCNSLFAGQSDLDQKCNVCHKPDRVERCMYE

YCLRGQQGFCDHAWEFKKECYIKHGDTLEVPDECKGSLLGGPSVFLFPPKPKDTLMISRTPEVTCVVVDVSQE

DPDVKFNWYVNGAEVHHAQTKPRETQYNSTYRVVSVLTVTHQDWLNGKEYTCKVSNKALPAPIQKTISKDKG

QPREPQVYTLPPSREELTKNQVSLTCLVKGFYPSDIVVEWESSGQPENTYKTTPPVLDSDGSYFLYSKLTVDKS

RWQQGNVFSCSVMHEALHNHYTQKSLSVSPGK

### SUPPLEMENTAL FIGURES

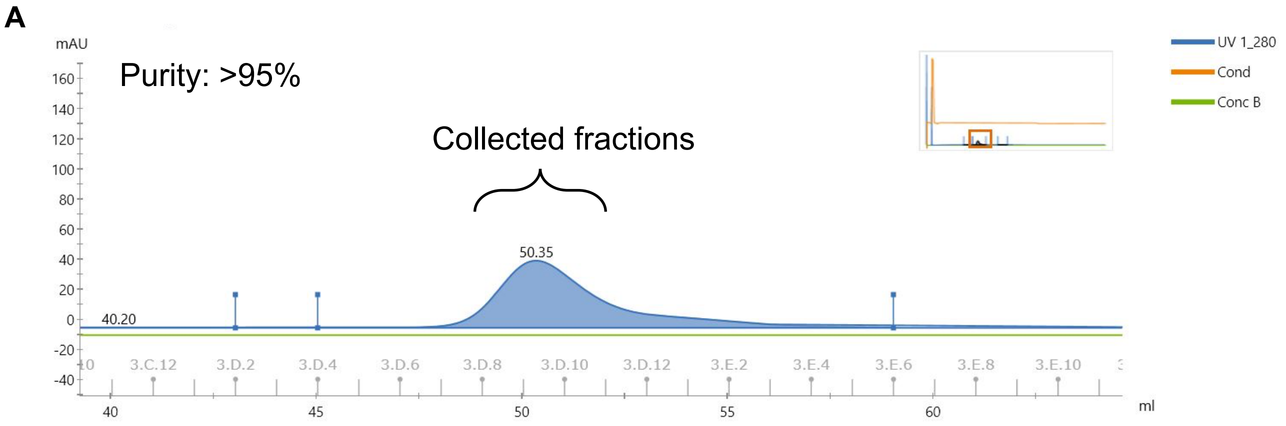


Peak Table - UV 1_280

| Peak | Retention ml | Area ml*mAU | Area % | Ext coeff. mg ml⁻¹ cm⁻¹ | Fraction(s) | Volume ml | Conductivity mS/cm |
|---|---|---|---|---|---|---|---|
| Peak A | 40.200 | 1.556 | 1.11 | | 3.C.6 - 3.D.1 | 8.000 | 16.64 |
| Peak B | 50.346 | 133.9 | 95.34 | | 3.D.4 - 3.E.5 | 13.999 | 16.62 |
| Peak C | 76.372 | 4.985 | 3.55 | | 3.F.7 - 3.G.4 | 10.002 | 16.56 |

**Figure S1. Purification of NHP-RMA proteins. B)** UV absorption profile of eluted NHP-RMA protein fractions obtained through size exclusion chromatography. Collected fractions achieved over 95% purity.

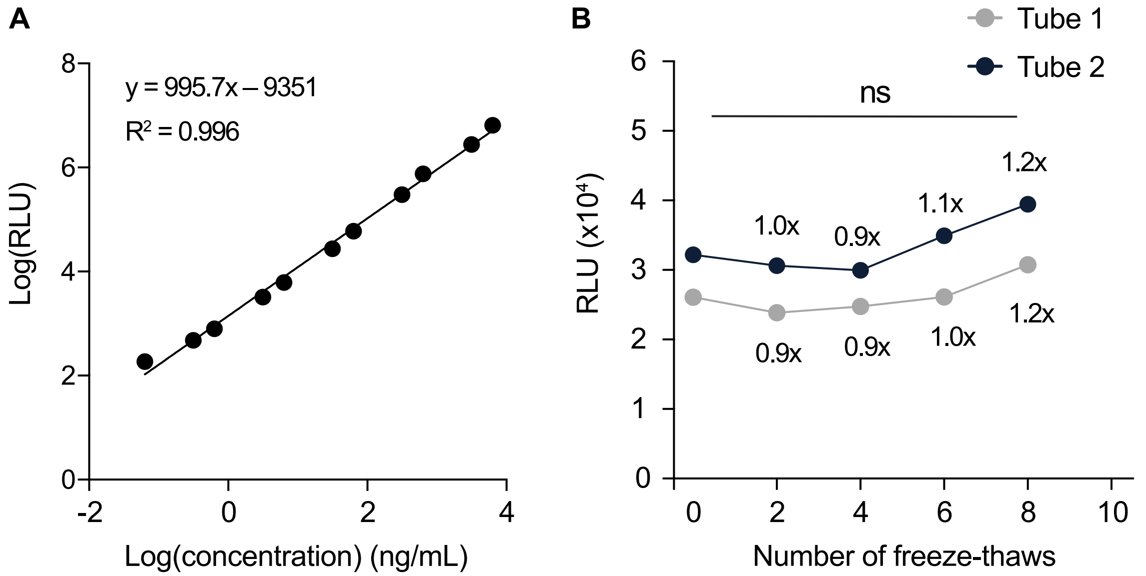


**Figure S2. NHP-RMA's standard curve and freeze-thaw effects. A) Standard curve of NHP-RMA proteins spiked into mouse serum, fitted using linear regression.** Bioluminescence signal values are presented as average of $n$ = 2 technical replicates. **B**) Luciferase activity of NHP-RMAs after repeated freeze thaw cycles. For each cycle, NHP-RMA proteins spiked in mouse serum were frozen at -80°C for 1 to 24 hrs, then thawed at room temperature before measuring their luciferase activity. Numbers represent signal fold increases relative to the baseline with no freeze-thaw cycles (i.e. 1x is equivalent to no change). One-way repeated measures ANOVA, with Dunnett's test (all $p > 0.05$).

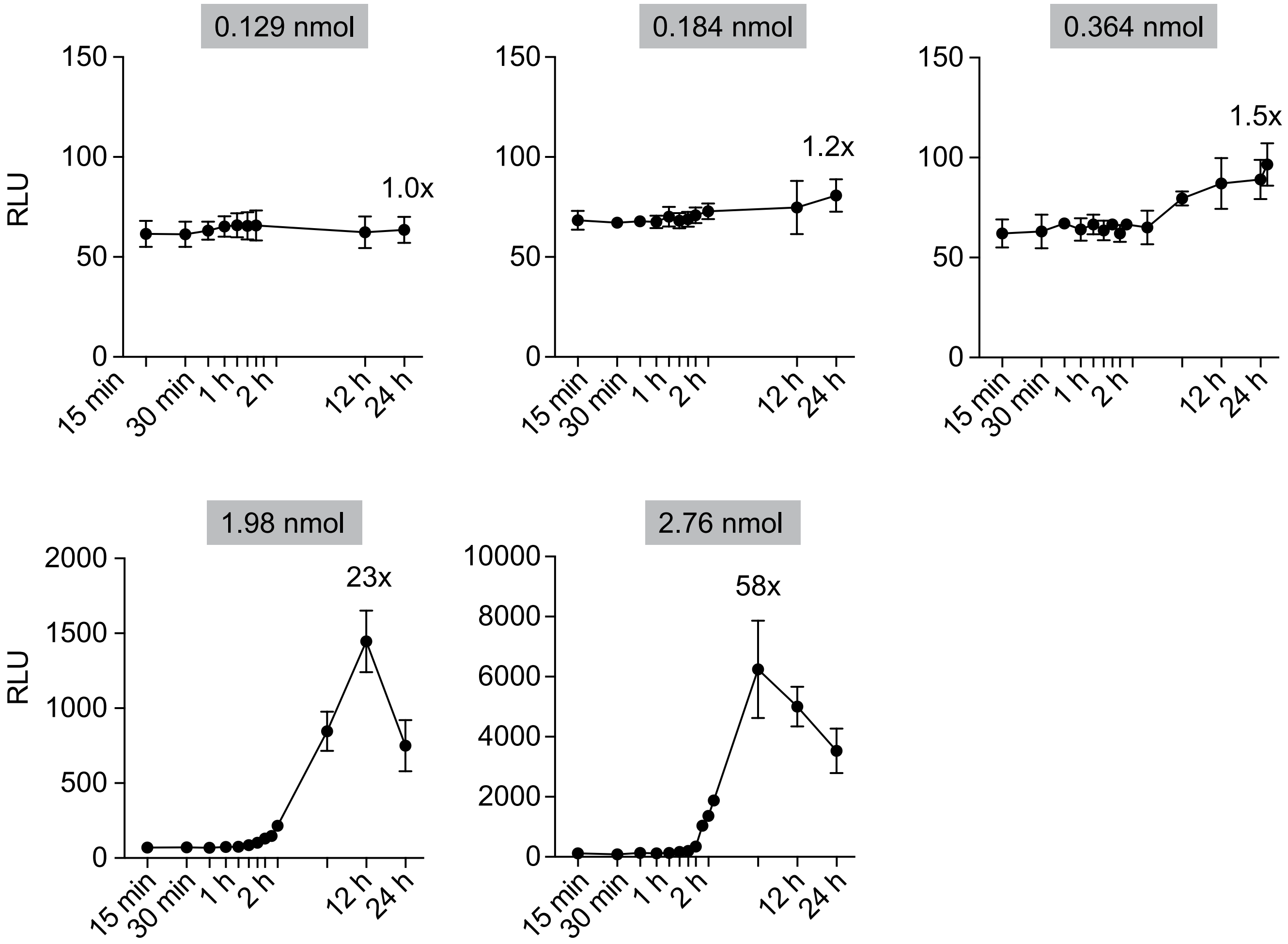


**Figure S3. Bioluminescence signals in serum samples following direct NHP-RMA protein injection.** Same data as in **Fig. 2c**, showing individual macaque NHP-RMA protein signals as a function of time for different injection doses. Numbers indicate signal fold increases relative to the pre-injection baseline signal. The number of technical replicates (*n* = 2; 0.364, 1.98, and 2.76 nmol doses) and 8 (*n* = 14; 0.129 and 0.184 nmol doses) for serum luciferase assays of the same blood samples. Data are shown as mean ± S.D.

#### SUPPLEMENTAL TABLES

| ID | Sex | Age | Weight | Intravenous (µg/kg) | Protein (nmol) | AAV (Transgene) |
|---|---|---|---|---|---|---|
| B1D0 | M | 8.1 | 10.3 | 329.16 | | |
| A35C | M | 12.2 | 13.1 | 1.60 | | |
| AB8A | F | 9.4 | 6.2 | | 0.184 | |
| A985 | F | 9.7 | 5.9 | | 0.364; 2.78 | |
| A64B | F | 11.3 | 7.3 | | 1.98 | DIO-Gluc; DIO-hM3dQ; Cre; Cluc |
| AAED | F | 11.8 | 9.1 | | 0.129 | DIO-Gluc; DIO-hM3dQ; Cre; Cluc |
| B00E | F | 9.0 | 6.4 | | | Gluc; Cluc |
| ABB4 | F | 9.8 | 11.85 | | | Gluc; Cluc |
| C231 | M | 4.1 | 8.4 | | | Gluc; Cluc |

**Table S1. Nonhuman primate demographics and experimental assignment.**

| | | | | | | | | | | | Blood Draws (1.5 mL) | | | |
|---|---|---|---|---|---|---|---|---|---|---|---|---|---|---|
| | | | | | | | | | Rate (µL/min) | | Early | Late | | |
| ID | Construct | Luciferase | nmol | µg | Site | Hemi | µL | Injections | Start | End | Minutes | Hours | | |
| AAED | Protein | Gluc | 0.129 | 4.05 | Put | L | 40.50 | 1 | 0.5 | 3.0 | 0:15:150 | | 12 | 24 |
| | | | | | Put | R | 40.50 | 1 | 0.5 | 3.0 | | | | |
| AB8A | Protein | Gluc | 0.184 | 4.58 | Put | L | 57.20 | 1 | 0.5 | 3.0 | 0:15:105 | | 12 | 24 |
| A985 | Protein | Gluc | 0.364 | 15.95 | Put | R | 24.95 | 2 | 0.5 | 3.0 | 0:15:195 | 6 | 12 | 24 |
| A64B | Protein | Gluc | 1.980 | 41.43 | Put | L | 94.23 | 2 | 0.5 | 3.0 | 0:15:165 | 6 | 12 | 24 |
| | | | | | Put | R | 92.82 | 2 | 0.5 | 3.0 | | | | |
| A985 | Protein | Gluc | 2.760 | 43.85 | Put | L | 99.00 | 2 | 0.5 | 3.0 | 0:15:160 | 6 | 12 | 24 |
| | | | | | Put | R | 99.00 | 2 | 0.5 | 3.0 | | | | |

**Table S2. NHP-RMA Protein Injection (pSL18.1) and Blood Draw Details.** L = Left hemisphere; R = Right hemisphere; Put = Putamen

| | | | | | | | | | Rate (μL / min) | | Blood Draws (1.5 mL) | | |
|---|---|---|---|---|---|---|---|---|---|---|---|---|---|
| **ID** | **AAV** | **pM** | **Transgene** | **Titer** | **Site** | **Hemi** | **μL** | **vg** | **Start** | **End** | **Minutes** | **Hours** | **Weeks** |
| B00E | AAV2 | hSyn | Gluc-IRES-GFP | 1E+12 | NAc | L | 9.04 | 9.04E+12 | 0.5 | 3.5 | 0 | 24 | 1:1:11 |
| | AAV2 | hSyn | Gluc-IRES-GFP | 1E+12 | NAc | R | 9.02 | 9.02E+12 | 0.5 | 3.5 | | | |
| | AAV2 | hSyn | Gluc-IRES-GFP | 1E+12 | 32 | L | 3.77 | 3.77E+12 | 0.2 | 1.5 | | | |
| | AAV2 | hSyn | Gluc-IRES-GFP | 1E+12 | 32 | R | 3.92 | 3.92E+12 | 0.2 | 1.5 | | | |
| | AAV2 | hSyn | Gluc-IRES-GFP | 1E+12 | 9/10 | L | 3.91 | 3.91E+12 | 0.2 | 1.5 | | | |
| | AAV2 | hSyn | Gluc-IRES-GFP | 1E+12 | 9/10 | R | 5.58 | 5.58E+12 | 0.2 | 1.5 | | | |
| | AAV2 | hSyn | Cluc | 1E+12 | NAc | L | 9.04 | 9.04E+12 | 0.5 | 3.5 | | | |
| | AAV2 | hSyn | Cluc | 1E+12 | NAc | R | 9.02 | 9.02E+12 | 0.5 | 3.5 | | | |
| | AAV2 | hSyn | Cluc | 1E+12 | 32 | L | 3.77 | 3.77E+12 | 0.2 | 1.5 | | | |
| | AAV2 | hSyn | Cluc | 1E+12 | 32 | R | 3.92 | 3.92E+12 | 0.2 | 1.5 | | | |
| | AAV2 | hSyn | Cluc | 4E+12 | 9/10 | L | 3.91 | 1.56E+13 | 0.2 | 1.5 | | | |
| | AAV2 | hSyn | Cluc | 4E+12 | 9/10 | R | 5.58 | 2.23E+13 | 0.2 | 1.5 | | | |
| ABB4 | AAV2 | hSyn | Gluc-IRES-GFP | 1E+12 | Put | L | 86.55 | 8.66E+13 | 0.5 | 3.5 | 0:30:120 | 24 | 1:1:6 |
| | AAV2 | hSyn | Cluc | 1E+12 | Put | R | 90.86 | 9.09E+13 | 0.5 | 3.5 | | | |
| C231 | AAV2 | hSyn | Cluc | 1E+12 | Put | L | 88.50 | 8.85E+13 | 0.5 | 3.5 | 0:30:120 | 24 | 1:1:8 |
| | AAV2 | hSyn | Gluc-IRES-GFP | 1E+12 | Put | R | 90.98 | 9.10E+13 | 0.5 | 3.5 | | | |
| A64B | AAV2 | hSyn | Cluc | 1E+12 | NAc | R | 23.29 | 2.33E+13 | 0.5 | 3.5 | 0:30:120 | 24 | 1:1:10 |
| | AAV2 | hSyn | CRE-eGFP | 1E+12 | NAc | R | 72.21 | 7.22E+13 | 0.5 | 3.5 | | | |
| | AAV2 | hSyn | DIO-Gluc-IRES-GFP | 1E+12 | BLA | R | 26.36 | 2.64E+13 | 0.5 | 2.0 | | | |
| | AAV2 | hSyn | DIO-Gluc-IRES-GFP | 4E+12 | BLA | R | 25.04 | 1.00E+14 | 0.5 | 2.0 | | | |
| | AAV2 | hSyn | DIO-hM3Dq-mCherry | 1E+12 | BLA | R | 26.36 | 2.64E+13 | 0.5 | 2.0 | | | |
| AAED | AAV2 | hSyn | Cluc | 1E+12 | NAc | L | 23.85 | 2.39E+13 | 0.5 | 3.5 | 0:30:120 | 24 | 1:1:10 |
| | AAV2 | hSyn | CRE-eGFP | 1E+12 | NAc | L | 73.92 | 7.39E+13 | 0.5 | 3.5 | | | |
| | AAV2 | hSyn | DIO-Gluc-IRES-GFP | 1E+12 | BLA | L | 48.11 | 4.81E+13 | 0.5 | 2.0 | | | |
| | AAV2 | hSyn | DIO-Gluc-IRES-GFP | 4E+12 | BLA | L | 24.80 | 9.92E+13 | 0.5 | 2.0 | | | |
| | AAV2 | hSyn | DIO-hM3Dq-mCherry | 1E+12 | BLA | L | 48.11 | 4.81E+13 | 0.5 | 2.0 | | | |
| | AAV2 | hSyn | Cluc | 1E+12 | 9/10 | R | 10.45 | 1.05E+13 | 0.2 | 1.5 | | | |

**Table S3. NHP-RMA AAV Injections and Blood Draw Details.** pM = Promotor; L = Left hemisphere; R = Right hemisphere; Put = Putamen; BLA = Basolateral amygdala; NAc = Nucleus accumbens; 9/10 = Area 9/10 (frontopolar cortex); 32 = Area 32 (anterior cingulate cortex).
